# Evolution and development of male-specific leg brushes in Drosophilidae

**DOI:** 10.1101/786764

**Authors:** Kohtaro Tanaka, Olga Barmina, Ammon Thompson, Jonathan H. Massey, Bernard Y. Kim, Anton Suvorov, Artyom Kopp

## Abstract

The origin, diversification, and secondary loss of sexually dimorphic characters are common in animal evolution. In some cases, structurally and functionally similar traits have evolved independently in multiple lineages. Prominent examples of such traits include the male-specific grasping structures that develop on the front legs of many dipteran insects. In this report, we describe the evolution and development of one of these structures, the male-specific “sex brush”. The sex brush is composed of densely packed, irregularly arranged modified bristles and is found in several distantly related lineages in the family Drosophilidae. Phylogenetic analysis using 250 genes from over 200 species provides modest support for a single origin of the sex brush followed by many secondary losses; however, independent origins of the sex brush cannot be ruled out completely. We show that sex brushes develop in very similar ways in all brush-bearing lineages. The dense packing of brush hairs is explained by the specification of bristle precursor cells at a near-maximum density permitted by the lateral inhibition mechanism, as well as by the reduced size of the surrounding epithelial cells. In contrast to the female and the ancestral male condition, where bristles are arranged in stereotypical, precisely spaced rows, cell migration does not contribute appreciably to the formation of the sex brush. The complex phylogenetic history of the sex brush can make it a valuable model for investigating coevolution of sex-specific morphology and mating behavior.

## Introduction

Most animals are sexually dimorphic. Perhaps the most fascinating feature of sexual dimorphism is the rapid evolutionary turnover of sex-specific traits. Even among close relatives, the characters that distinguish males from females vary greatly from species to species. This simple observation implies that new sexual characters are gained, and ancestral ones are often lost, during the evolution of many if not most animal lineages. Understanding the genetic and developmental basis of this turnover is necessary to shed light on one of the most important drivers of biological diversity.

Most higher Diptera mate with the male on top of the female, and the male front (T1) legs are often involved in grasping or stimulating the female (Huber et al., 2007; McAlpine, 1981). Perhaps for this reason, male-specific ornaments or grasping structures are found on the T1 legs of many dipteran species (Daugeron et al., 2011; Eberhard, 2001; Ingram et al., 2008; Sivinski, 1997). In Drosophilidae, two of the most obvious male-specific leg modifications include the sex combs found in the *Drosophila melanogaster* and *obscura* species groups of subgenus *Sophophora* and in the genus *Lordiphosa* (Katoh et al., 2018; Kopp, 2011); and tarsal sex brushes, which are found in several lineages and are the focus of this study (Figure 1). Both types of structures are composed of modified mechanosensory bristles and develop at the same location in the T1 leg, namely on the anterior-ventral surface of proximal tarsal segments. In females, as well as in males of sexually monomorphic species, this area is occupied by transverse bristle rows (TBR) – tightly packed parallel rows of mechanosensory bristles arranged perpendicular to the proximo-distal leg axis, which the flies use to clean their head (Kopp, 2011) (Figure 1 M, N). Due to this stereotypical positioning, it is straightforward to establish the homology of precursor tissues between males and females as well as across different Drosophilid lineages.

**Figure 1.**
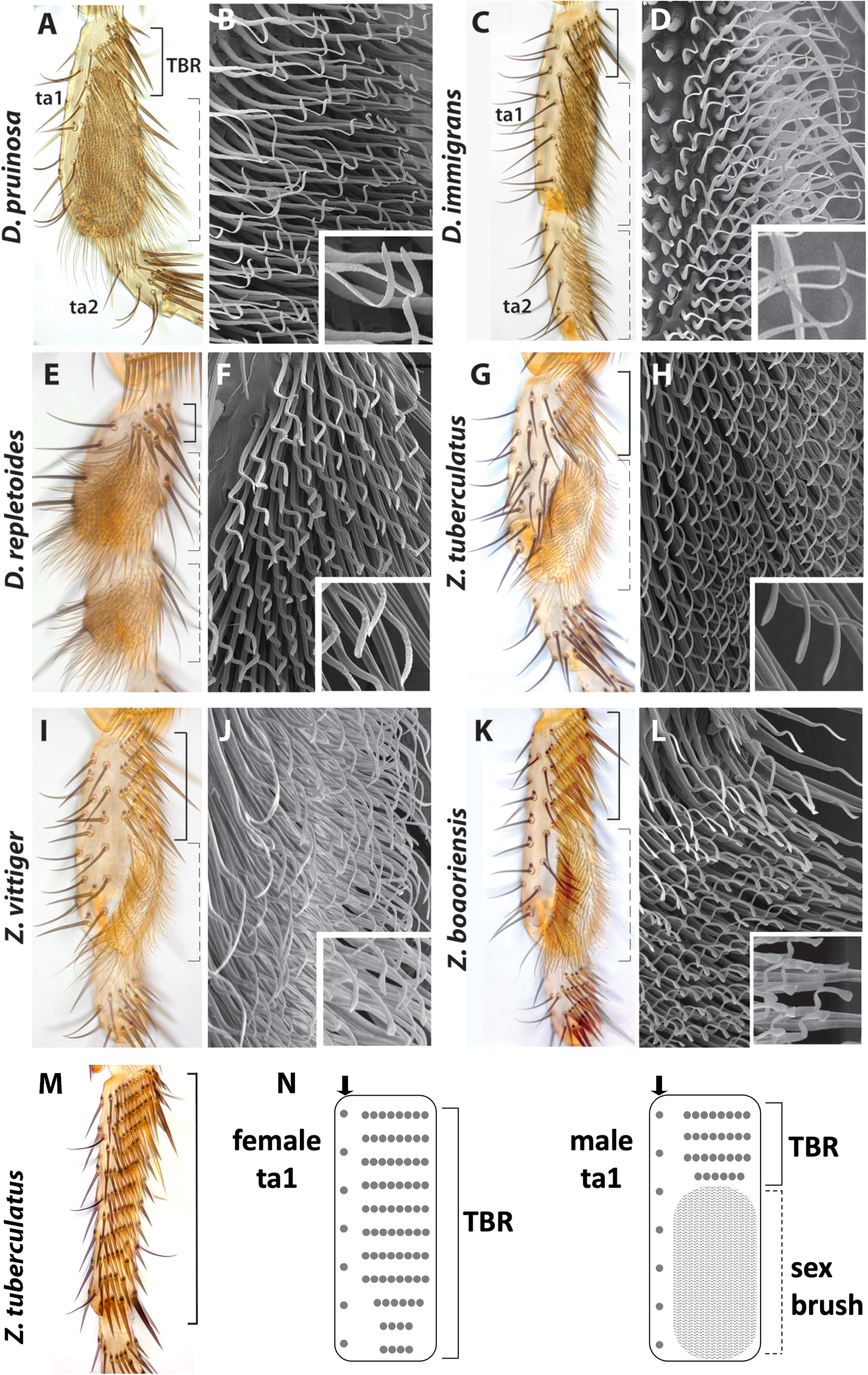
Sex brush morphology in distantly related species. (A, C, E, G, I, K) Brightfield images of the first and second tarsal segments (ta1 and ta2) of the front leg in males of six species. (B, D, F, H, J, L) SEM images of the ta1 sex brush. A-B) *D. pruinosa*. A) The brush (shown by dotted brackets in this and other panels) occupies the distal ~80% of ta1, replacing most of the transverse bristle rows (TBRs, shown by solid brackets), which in females cover the entire anterior-ventral surface. The ta1 segment is slightly widened at the distal end. B) The tips of brush hairs are flattened, pointed, and form hooks that curve toward the base of the leg (see inset). C-D) *D. immigrans*. C) The brush covers the distal ~60% of ta1 and most of ta2. The shape of the segments is not modified. D) Similar to *D. pruinosa*, the brush hairs are flattened, pointed, and form proximally curving hooks at the tips (inset). E-F) *D. repletoides*. E) The brush covers ~70% of ta1 and most of ta2. Both segments are shortened and have a bulbous shape. F) The tips of brush hairs are flattened but thick and blunt. They form hooks that curve away from the leg base (inset). G-H) *Z. tuberculatus*. G) The brush covers the distal ~60% of ta1. H) The tips of brush hairs curve distally and have a pointed paddle-like shape with a slight depression (inset). I-J) *Z. vittiger*. This species has sex brush very similar to *Z. tuberculatus*. K-L) *Z. bogoriensis*. Most hairs have tips that curve slightly toward the distal end of the leg. M) Brightfield image of the first tarsal segment of the front leg in the female *Z. tuberculatus*. TBRs occupy the entire anterior ventral surface. N) Schematic diagrams of the first tarsal segment of the female and male *Z. tuberculatus* showing the difference in bristle patterns. Arrow, longitudinal bristle row.

Previous studies have shown that similar sex comb morphologies have evolved multiple times in *Sophophora* and *Lordiphosa* (Kopp, 2011). Remarkably, this phenotypic convergence conceals divergent developmental mechanisms, as different cellular processes can produce similar adult structures. In some species, sex comb development relies on sex-specific patterning of bristle precursor cells, while in others it occurs through male-specific migration of sexually monomorphic precursors (Atallah et al. 2009; Tanaka et al. 2009; Atallah et al 2012).

The presence of sex brushes in multiple species provides another model for testing whether different cellular mechanisms can give rise to similar adult morphologies. Sex brushes are found in at least four groups within Drosophilidae: the *Drosophila immigrans* species group, the *loiciana* species complex, *D. repletoides*, and the genus *Zaprionus* (Table 1). In the *immigrans* group (see Supplemental Text for a discussion of this taxon), male sex brushes are found in some but not all of the species; the most likely scenario is that the brush was present in the last common ancestor of this clade, but was secondarily lost in the *nasuta* subgroup and greatly reduced in several other species (Rice et al., 2018). In *Zaprionus*, sex brushes are present in most African species, with the exception of three distantly related species that likely reflect independent secondary losses (Tsacas and Chassagnard, 1990; Yassin et al., 2008; Yassin and David, 2010). The Oriental *Anaprionus* subgenus of *Zaprionus* also contains species that lack sex brushes; however, this likely reflects the polyphyly of *Anaprionus* (see Supplemental Text). In any case, the sex brush was present at or near the base of *Zaprionus*. In the *loiciana* species complex, all described species have male sex brushes (Tsacas, 2002; Tsacas and Chassagnard, 2000). Finally, the fourth lineage where a sex brush is found consists of a single species, *D. repletoides* (See Supplemental Text and Table 1 for additional details on the taxonomic affiliation and leg morphology of the species used in this study).

**Table 1.**
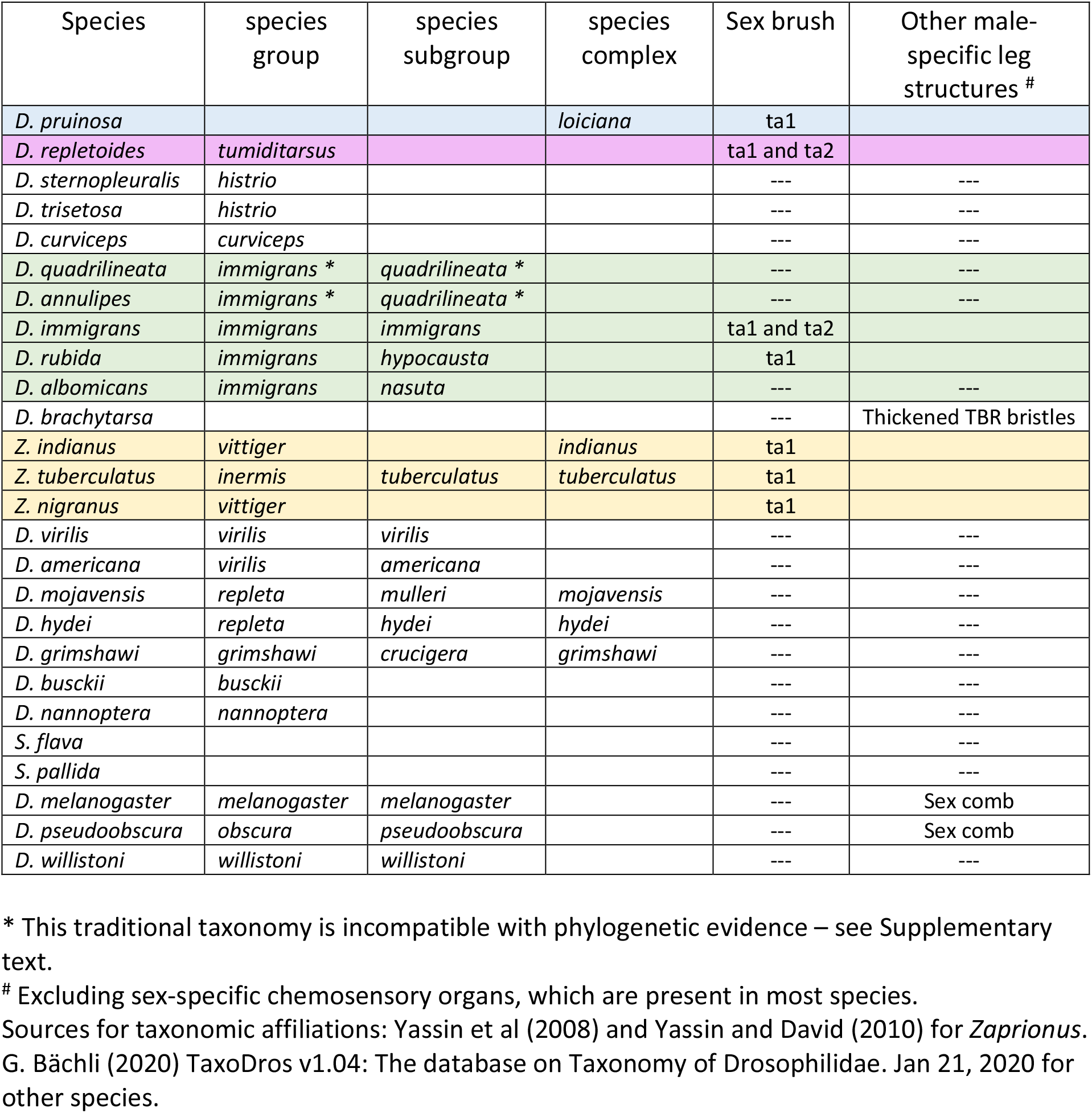
Taxonomic affiliations and notes on the leg morphology of brush-bearing species and their brush-less relatives used in this study.

| Species | species group | species subgroup | species complex | Sex brush | Other male-specific leg structures # |
| --- | --- | --- | --- | --- | --- |
| *D. pruinosa* |  |  | *loiciana* | ta1 |  |
| *D. repletoides* | *tumiditarsus* |  |  | ta1 and ta2 |  |
| *D. sternopleuralis* | *histrio* |  |  | --- | --- |
| *D. trisetosa* | *histrio* |  |  | --- | --- |
| *D. curviceps* | *curviceps* |  |  | --- | --- |
| *D. quadrilineata* | *immigrans* * | *quadrilineata* * |  | --- | --- |
| *D. annulipes* | *immigrans* * | *quadrilineata* * |  | --- | --- |
| *D. immigrans* | *immigrans* | *immigrans* |  | ta1 and ta2 |  |
| *D. rubida* | *immigrans* | *hypocausta* |  | ta1 |  |
| *D. albomicans* | *immigrans* | *nasuta* |  | --- | --- |
| *D. brachytarsa* |  |  |  | --- | Thickened TBR bristles |
| *Z. indianus* | *vittiger* |  | *indianus* | ta1 |  |
| *Z. tuberculatus* | *inermis* | *tuberculatus* | *tuberculatus* | ta1 |  |
| *Z. nigranus* | *vittiger* |  |  | ta1 |  |
| *D. virilis* | *virilis* | *virilis* |  | --- | --- |
| *D. americana* | *virilis* | *americana* |  | --- | --- |
| *D. mojavensis* | *repleta* | *mulleri* | *mojavensis* | --- | --- |
| *D. hydei* | *repleta* | *hydei* | *hydei* | --- | --- |
| *D. grimshawi* | *grimshawi* | *crucigera* | *grimshawi* | --- | --- |
| *D. busckii* | *busckii* |  |  | --- | --- |
| *D. nannoptera* | *nannoptera* |  |  | --- | --- |
| *S. flava* |  |  |  | --- | --- |
| *S. pallida* |  |  |  | --- | --- |
| *D. melanogaster* | *melanogaster* | *melanogaster* |  | --- | Sex comb |
| *D. pseudoobscura* | *obscura* | *pseudoobscura* |  | --- | Sex comb |
| *D. willistoni* | *willistoni* | *willistoni* |  | --- | --- |

\* This traditional taxonomy is incompatible with phylogenetic evidence – see Supplementary text.

The four clades of interest – *Zaprionus*, the *immigrans* species group, *D. repletoides*, and the *loiciana* complex – have never been included together in the same molecular phylogeny. Different combinations of these taxa have been examined in several studies, which were based on a small number of loci and produced conflicting results. Da Lage et al (Da Lage et al., 2007) and Yassin et al (Yassin et al., 2010) provided some evidence for a distant relationship among *Zaprionus*, *D. repletoides*, and the *immigrans* species group. Russo et al (Russo et al., 2013) placed *D. pruinosa* as sister to *D. sternopleuralis* (a member of the *histrio* species group, which lacks a sex brush), and the resulting clade as sister to the *immigrans* species group. The phylogenies of Da Lage et al (Da Lage et al., 2007) and Izumitani et al (Izumitani et al., 2016) did not include *D. pruinosa*, but did not support a sister-group relationship between *D. sternopleuralis* and the *immigrans* species group.

In this study, we used larger multilocus datasets to test whether the male sex brush evolved independently in each of these four clades, or whether its distribution could be better explained by a shared origin. In parallel, we compared the cellular mechanisms that produce the sex brush in different species. Our results show that the sex brushes of distantly related species develop through virtually identical developmental process. However, it remains unclear whether this similarity reflects a single origin or convergent evolution.

## Materials and Methods

### Leg imaging

For brightfield imaging, male front legs were dissected, mounted in Hoyers media between two coverslips, and photographed on a Leica DM500B microscope with a Leica DC500 camera. For electron microscopy, adult legs were dehydrated in 100% ethanol, critical point dried, and coated with gold. Scanning electron micrographs were taken on Thermo Fisher Quattro S and Philips XL30 TMP. Three to five legs were imaged for each species.

### Sequence data collection

The sources of live *Drosophila* strains and fixed specimens used in this study are listed in Supplemental Table S1. DNA was extracted from live or alcohol-fixed flies using an affinity resin-based protocol (Hi Yield^®^ Genomic DNA Mini Kit, Süd-Laborbedarf Gauting, Germany). PCR was carried out using DreamTaq polymerase (Thermo Fisher) and the following cycling conditions: 95° 5’ => (95° 30” => 55° 30” => 72° 80”) x 35 => 72° 5’ =>12°; the loci and primer sequences are listed in Supplemental Table S2. In some cases, two rounds of PCR with nested primers were needed to obtain amplicons from old, ethanol-fixed specimens. Amplified fragments were gel-purified and sequenced from both ends using amplification primers. Sequence chromatograms were trimmed in SnapGene Viewer, and the two end reads were aligned and edited in Geneious (Kearse et al., 2012). Heterozygous nucleotide positions, if present, were represented by IUPAC ambiguity codes. All new sequences were deposited in Genbank under accession numbers listed in Supplemental Table S1. Additional sequences were obtained from Genbank or extracted from whole-genome assemblies using Blast v2.2.23 (Supplemental Table S1).

### Phylogenetic tree reconstruction

We conducted two separate phylogenetic analyses. In the first, we used eight loci sequenced above (Supplemental Table S1) to reconstruct the phylogeny of selected representatives of brush-bearing clades and their closest brush-less relatives indicated by previous phylogenetic and taxonomic studies. The sequences of each locus were aligned using the MUSCLE algorithm (Edgar, 2004) implemented in Geneious (Kearse et al., 2012). The alignments were trimmed at the ends, and poorly aligning intronic regions were removed. The alignments of all eight loci were then concatenated for combined analysis. Combined Bayesian analysis was carried out in MrBayes v3.2.6 (Ronquist et al., 2012). Two sets of analyses were conducted. In the first, the dataset was partitioned by gene, and each locus was allowed to follow a different nucleotide substitution model with empirically estimated parameters. In the second, we partitioned the dataset by gene and by codon position, and used PartitionFinder with the PhyML algorithm (Guindon et al., 2010; Lanfear et al., 2012; Lanfear et al., 2017) to identify the appropriate partitioning scheme; this resulted in a total of 21 character subsets, each of which was subsequently allowed to follow its own substitution model with empirically estimated parameters. For each analysis, two parallel runs of 1,500,000 generations, each starting from a different random tree, were carried out, and convergence was confirmed by comparing tree likelihoods and model parameters between the two runs. *D. melanogaster* was used as outgroup. Trees were sampled every 1000 generations and summarized after a 20% relative burn-in. Samples of probable trees were extracted from tre.tprobs file, and a strict consensus of most probable trees with combined posterior probabilities of 95% or 99% was constructed from each set of trees in Geneious. Consensus trees were formatted using FigTree v1.3.1 (http://tree.bio.ed.ac.uk/software/figtree/).

In the second analysis, we reconstructed the phylogeny of >200 species of Drosophilidae using 250 single-copy BUSCO genes extracted from complete genome assemblies (Supplemental Table S3). Several important species, including *D. trisetosa*, *D. sternopleuralis*, and *D. brachytarsa* could not be included in this analysis since their genome sequences were not available. First, we estimated multiple sequence alignments (MSAs) using the L-INS-I strategy in MAFFT v7.453 (Katoh and Standley, 2013). Sites containing less than 3 non-gap characters were removed. After trimming, the MSA lengths ranged from 325 bp to 16,465 bp with an average length of 2,289 bp. Next, for phylogenetic inference we concatenated 250 MSAs to form a supermatrix containing 572,343 sites in total. We used this supermatrix as a single partition to estimate a maximum-likelihood (ML) tree topology in IQ-TREE v1.6.5 (Nguyen et al., 2015), specifying the GTR+I+G substitution model (Supplemental Figure S4). To estimate the support for each node in the resulting topology, we computed three different reliability measures. We did 1,000 ultrafast bootstrap (UFBoot) replicates (Minh et al., 2013), an approximate likelihood ratio test with the nonparametric Shimodaira-Hasegawa correction (SH-aLRT), and a Bayesian-like transformation of aLRT (Anisimova et al., 2011).

We implemented the Bayesian algorithm of MCMCTree v4.9h (Yang, 2007) with approximate likelihood computation to estimate divergence times using five fossils for age prior construction, analogous to the calibration scheme A described in (Suvorov et al., 2022). First, we divided our 250 loci into five equal datasets and generated five supermatrices consisting of 50 MSAs each. We used these datasets to perform the dating analyses. For each of our five datasets, we estimated branch lengths by ML and then the gradient and Hessian matrices around these ML estimates in MCMCTree using the DNA supermatrix and species tree topology estimated by IQ-TREE. Then, we used the gradient and Hessian matrix, which constructs an approximate likelihood function by Taylor expansion (Reis and Yang, 2011), to perform fossil calibration in an MCMC framework. For this step, we specified a GTR+G substitution model with four gamma categories; birth, death and sampling parameters of 1, 1 and 0.1, respectively. To model rate variation, we used an uncorrelated relaxed clock. To ensure convergence, the analysis was run three times independently for each of the 5 datasets for 7 × 10^6^ generations (first 2 ×10^6^ generations were discarded as burn-in), logging every 1,000 generations. Additionally, we performed sampling from the prior distribution only. Convergence was assessed for the combined MCMC runs for each of the 5 datasets using Effective Sample Size criteria (ESS > 100) in Tracer (Rambaut et al., 2018).

### Ancestral character reconstruction

We conducted ancestral character reconstruction using the 250-locus BUSCO dataset, in which >200 species were sampled without regard to the presence or absence of leg brushes. Our 8-locus dataset was not suitable for similar analysis since brush-bearing species were intentionally over-sampled in that dataset. To estimate the probability of the origin and secondary loss of sex brushes, we reconstructed ancestral character states using the time-calibrated BUSCO trees described above. Brush presence/ absence matrix was analyzed under five different models of trait evolution: (1) a simple MK model (Lewis, 2001) with unequal rates and a strict molecular clock; (2) an MK model with unequal rates and a random local relaxed clock (RLC; (Drummond and Suchard, 2010)); (3) a hidden states variable rates model with two latent rate classes (Beaulieu et al., 2013). The latter model assumes that each of the two binary character states (trait present vs trait absent) can exist in two discrete, not directly observable rate classes (“fast evolving / likely to change” vs “slow evolving / not likely to change”). We also used (4) an approximation of a Dollo model made by assuming that the rate of trait loss is more than 300 times greater than the rate of gain; and (5) a modified threshold model similar to Felsenstein’s (Felsenstein, 2005). This modified threshold model assumes that the gain of a trait proceeds sequentially through 9 ordered latent states, such that each state can only transition to its nearest neighbor state towards or away from the trait. This implies that a species that lacks the trait can exist in any of the 9 “trait-absent” states that are not directly observable. For species that lack sex brushes, we assume a uniform distribution over these latent states.

All models were implemented and analyzed in RevBayes (Höhna et al., 2016), and RevBayes MCMC outputs were analyzed in R using phytools (Revell, 2012). See Supplemental Table S4 for prior distribution assumptions for all model parameters. Under each of the above models, except RLC, and for each of the five trees inferred from different sets of genes, MCMC was run for 50,000 cycles with a sampling rate of 1 in 50 to produce at least 200 ESS for all parameters. Because the RLC model takes significantly longer to run, we ran the MCMC for a set amount of time instead of set number of cycles. This also achieved > 200 ESS for all RLC analyses. Two independent chains were run to confirm convergence to the same posterior. Sensitivity to prior specification was assessed by comparing the marginal posterior to marginal prior for each parameter. 100 stochastic character evolution histories were simulated during one of the MCMC chains for each of the models by sampling every tenth sample from the posterior distribution. The resulting simmap files of 100 character histories mapped to a fixed tree were then analyzed to infer the ancestral states at nodes and along branches as well as the number and type of transitions. For the RLC model, we checked sensitivity to the prior on the frequency of rate shifts by comparing inferences under a prior model with expected number of rate shifts of 2 and that of 10 and obtained very similar inferences.

### Immunocytochemistry and microscopy

Fly cultures were raised on standard *Drosophila* media at approximately 22°C. Since each species develops at a different rate, the timing of pupal stages was determined empirically based on the morphology of transverse bristle rows (TBRs) in the same leg. Each species was imaged at an early stage when TBR bristles of the tibia and the first tarsal segment are separated by one or more intervening epithelial cells; this stage corresponds to 16 - 21 hours after pupariation (AP) in *D. melanogaster*. Each species was also imaged at a late stage, after the packing of TBRs is completed and bristle shaft differentiation is underway (corresponding to 36+ hrs AP in *D. melanogaster*). Pupal legs were dissected, processed and immunostained as in Tanaka et al (2009). The primary antibodies used were rat anti-E-cadherin (DCAD2, from the Developmental Studies Hybridoma Bank, at 1:20) for *D. pruinosa* and *Z. tuberculatus*, and mouse anti-Armadillo (N2 7A1, DSHB; 1:30) for *D. immigrans* and *D. repletoides*. None of the tested antibodies had sufficient cross-reactivity in all examined species, forcing us to use different antibodies for different species. However, this was not an issue since our goal was to visualize the cell shape and not the subcellular structures. AlexaFluor 488 secondary antibodies (Invitrogen) were used at 1:400. Fluorescent images were taken on an Olympus 1000 confocal microscope. Images were processed using ImageJ and Adobe Photoshop. Legs from at least five individuals were examined per species per stage, and three to five legs were imaged. To assemble leg images, signals from non-epidermal tissues were removed from each confocal section, and Z-series projection was produced using Image J.

### Image analysis

To determine the ratio between bristle and epithelial cells, a group of 30 neighboring bristle cells was selected from the brush region of each leg processed as above. The epithelial cells associated with these bristle cells were manually counted. Cells on the periphery of the area were included if they shared a side with a bristle cell. For each stage and species, five legs were analyzed. 2-sample Wilcoxon rank-sum (a.k.a Mann-Whitney U test) function in R was used for statistical tests. For the analysis of epithelial cell size, cell outlines were manually traced using the polygon selection tool in Image J and cell areas were measured using the software’s ROI manager. 30 epithelial cells in the proximal TBR region and 30 in the middle of the brush were measured in each leg.

### Analysis of mating behavior

High-speed videos of mating behavior were recorded using a Fascam Photron SA4 mounted with a 105 mm AF Micro Nikkor lens. In brief, individual virgin males were isolated upon eclosion in food vials and aged for up to two weeks. Virgin females were isolated upon eclosion and housed in groups of 20-30. Pairs of males and females were then gently aspirated into single wells of a 96 well culture plate (Corning 05-539-200) filled halfway with a hardened 2% agarose solution and sealed using a glass microscope slide and tape. Video clips were captured at 1000 frames per second (fps) using Photron Fastcam Viewer software.

## Results

### Sex brush morphology is similar in distantly related species

Species descriptions found in taxonomic literature (see Supplemental text) report the presence of male-specific leg brushes but are often vague about their precise morphology and location. In all examined species of *Zaprionus*, the *immigrans* species group, *loiciana* species complex, and *D. repletoides*, we observed the sex brush on the anterior-ventral surface of the first (ta1) and sometimes also the second (ta2) tarsal segment of the front leg (Table 1). In females and in species that lack the sex brush, this area is occupied by transverse bristle rows (TBRs) (Figure 1M; Tokunaga 1962; Kopp 2011). The male-specific brush replaces the distal TBRs, with a few TBRs remaining at the proximal end of ta1 (Figure 1). In the *immigrans* group (Rice et al. 2018) and in the other species (Figure 1), we find several major differences between the brush and the TBRs. The bristles of each TBR are aligned into a straight, tightly packed row that is nearly perpendicular to the proximo-distal (PD) leg axis, while the consecutive TBRs along the PD axis are separated by many cell diameters (Figure 1M). In contrast, the modified bristles (“hairs”) that make up the brush are packed closely together in all directions and do not show any obvious regular arrangement. The shafts of the TBR bristles are robust and straight, with ridges and grooves running their length and a triangular bract at the base; the brush hairs are thin and wavy with a smooth surface and lack bracts (Figure 1).

To compare sex brush morphology among species at a finer scale, we imaged these structures using scanning electron microscopy (SEM). This analysis revealed a small but consistent difference. In *D. immigrans* and *D. pruinosa*, the tips of brush hairs are thin and flat, taper to a point, and form hooks that curve toward the base of the leg (Figure 1B, D). In *D. repletoides* the hair tips are noticeably thicker and curve away from the leg base (Figure 1 F). In all *Zaprionus* species examined, the hair tips are flattened into paddle-like shapes (Figure 1 H, J, L). Similar to *D. repletoides, Z. tuberculatus* and *Z. vittiger* have hair tips that curve distally (Figure 1 H, J), whereas in *Z. bogoriensis*, the direction of the curve is less pronounced. Thus, while the spatial arrangement of bristles appears to be similar in all species, subtle differences exist in the morphology of bristle shafts. These differences may reflect the phylogenetic relationships among these taxa, especially the close relationship between *D. pruinosa* and the *immigrans* species group (see next section).

### Single or multiple origins of male leg brushes?

Previous reports suggest that at least some of the brush-bearing lineages are more closely related to brush-less species than to other brush-bearing clades (Da Lage et al., 2007; Yassin et al., 2010; Russo et al., 2013; Izumitani et al., 2016). However, the four brush-bearing clades have never been included together in the same molecular phylogeny. To confirm these results, we used partial coding sequences of eight nuclear loci to reconstruct the phylogeny of selected representatives of brush-bearing clades and their closest brush-less relatives indicated by previous studies. Separate analyses of each locus produced very poorly resolved trees with polytomies at the basal nodes. We therefore combined the data from all eight loci (up to 9060 nucleotides per species) for a partitioned Bayesian analysis where each locus was allowed to follow its own empirically estimated substitution model, but all loci were constrained to the same tree topology. The resulting tree (Figure 2; brush-bearing species labeled in blue) suggests a close relationship of *D. pruinosa* to the *immigrans* species group, with the (*D. sternopleuralis* + *D. trisetosa*) clade, which belongs to the *histrio* species group, as the next outgroup. In contrast, *D. repletoides* is placed as sister group to the (*D. busckii* + *D. brachytarsa*) clade, well away from the *immigrans-pruinosa* lineage. Finally, the *Zaprionus* genus appears, based on this dataset, to be distantly related to both *D. repletoides* and the *immigrans-pruinosa* lineage. We also note that *D. curviceps* and *D. annulipes* appear as sister groups with 100% support, while there is no support for clustering the *immigrans* species group either with the (*D. curviceps* + *D. annulipes*) clade or with *D. quadrilineata* (see Supplemental text). A strict consensus of 11 trees with the cumulative posterior probability of 95% is not resolved near the base but does not support a close relationship among the brush-bearing lineages: *D. repletoides*, the *immigrans-pruinosa* clade, and *Zaprionus* (Figure S1). None of the 27 most probable trees, with the cumulative posterior probability of 99%, grouped *Zaprionus* with either *D. repletoides* or the *immigrans-pruinosa* clade, or the *repletoides-busckii*-*brachytarsa* clade with the *immigrans-pruinosa* clade. We repeated this analysis with a more complex partitioning scheme, where each locus and each codon position were allowed to follow their own substitution models. This analysis produced a tree with the same topology as in the simpler partitioning scheme, although with slightly different values of node support (Figure S2). In summary, we found substantial though not overwhelming support for a close relationship between *D. pruinosa*, and by implication the *loiciana* species complex, and the *immigrans* species group. However, based on this dataset, the probability of a close relationship among the different brush-bearing clades – *D. repletoides*, *Zaprionus*, and the *immigrans-pruinosa* lineage – is low for each of the possible pairwise relationships.

**Figure 2.**
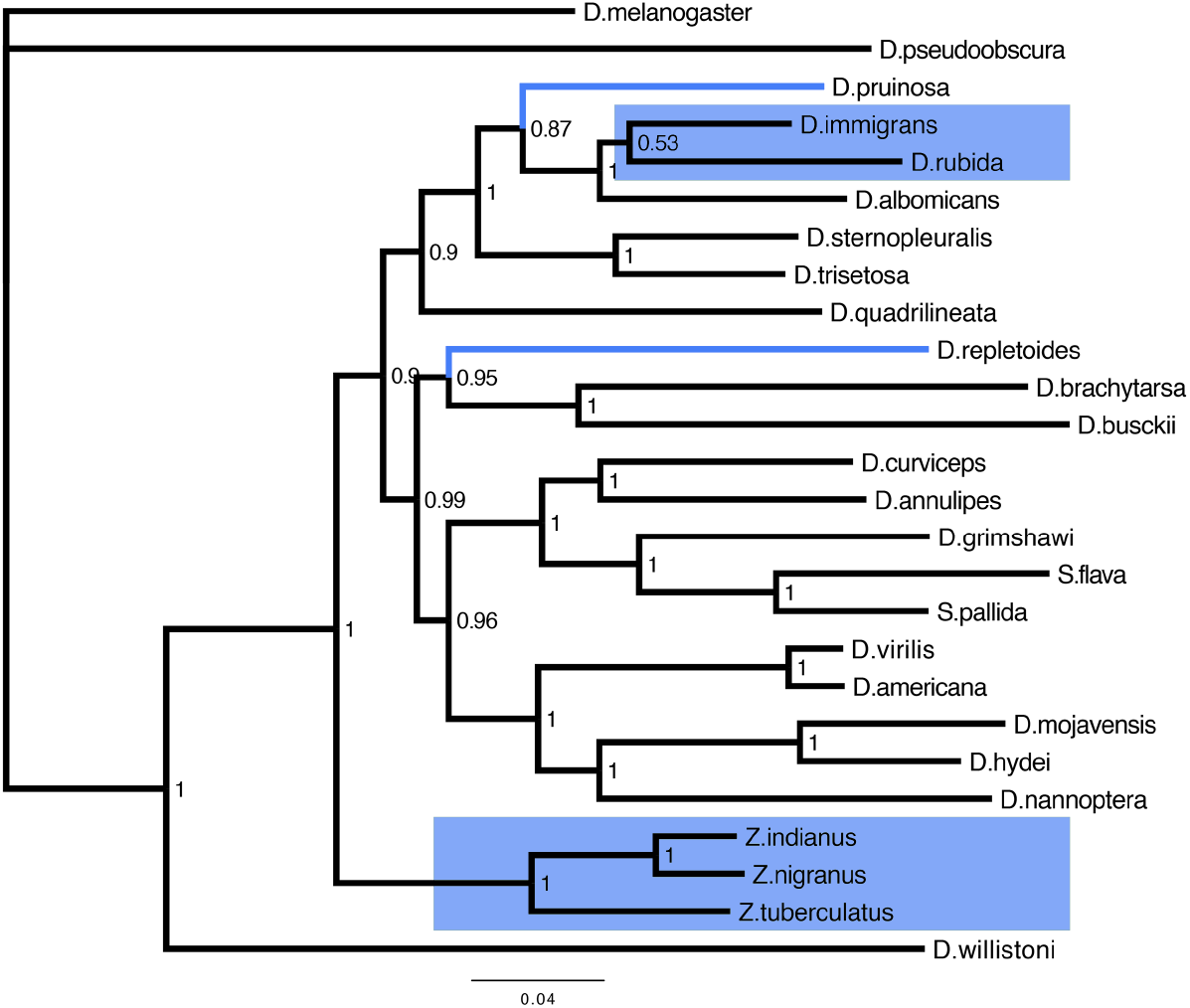
Brush-bearing groups are more closely related to brush-less species than to each other. This Bayesian phylogeny is based on the combined 8-locus dataset. Numbers at each node indicate the posterior probabilities of the respective taxon bipartitions. Species and clades with male sex brushes are highlighted in blue.

This finding has two potential explanations for the observed distribution of brush-bearing groups. First, the leg brush may have evolved independently in each of the three (or four) brush-bearing clades. Alternatively, the brush may have evolved once in a common ancestor of all these lineages but was subsequently lost in some of its descendants. The 8-locus dataset is not suitable for distinguishing between these scenarios, because of its intentionally biased taxon sampling. To estimate the probability of the single-gain *vs* multiple-gains models of sex brush evolution, we used the larger, 250 gene / >200 species BUSCO dataset in which species were sampled with no regard for the presence or absence of leg brushes. In this taxon sample, the smallest clade that encompasses all four brush-bearing groups also contains many species that lack sex brushes (Figure 3, Figure S3, Figure S4). Under most models of character evolution, ancestral character reconstruction suggests that the sex brush originated once in the most recent common ancestor of the *immigrans* species group, *D. pruinosa*, *Zaprionus*, and *D. repletoides*. The probability that this common ancestor had a sex brush is 0.64 under the hidden states variable rates model (Beaulieu et al., 2013), and 0.77-0.87 under the MK models (Drummond and Suchard, 2010; Lewis, 2001) (Figure 3; Figure S4 A-C). As expected, enforcing an approximate Dollo model by assuming a highly informative prior where the rate of trait loss is more than 300 times higher than the rate of trait gain leads to a stronger conclusion (1.00 probability of single origin). However, this model also infers a ~0.5 probability that the last common ancestor of all Drosophilidae had a sex brush (Figure S4E). We believe this scenario is unlikely, and that this inference may be driven in part by under-sampling of basal drosophilid lineages in the BUSCO dataset. Another outlier result in the ancestral character reconstruction is produced by the threshold model with latent ordered states, which accommodates more gradual transitions between the “present” and “absent” states of discrete characters. This model puts the probability that the most recent common ancestor of the *immigrans* species group, *D. pruinosa, Zaprionus*, and *D. repletoides* had a sex brush at 0.17 (Figure S4D), thus favoring multiple independent gains of these structures. We note that all these analyses are likely to be biased in favor of inferring a single origin, because taxon sampling in the BUSCO phylogeny is very sparse for the *cardini, guarani, quinaria, guttifera, pallidipennis*, and other brush-less species groups that fall within the large *immigrans-repletoides* clade (Figure 3, Figure S3) (Kim et al., 2021; Suvorov et al., 2022).

**Figure 3.**
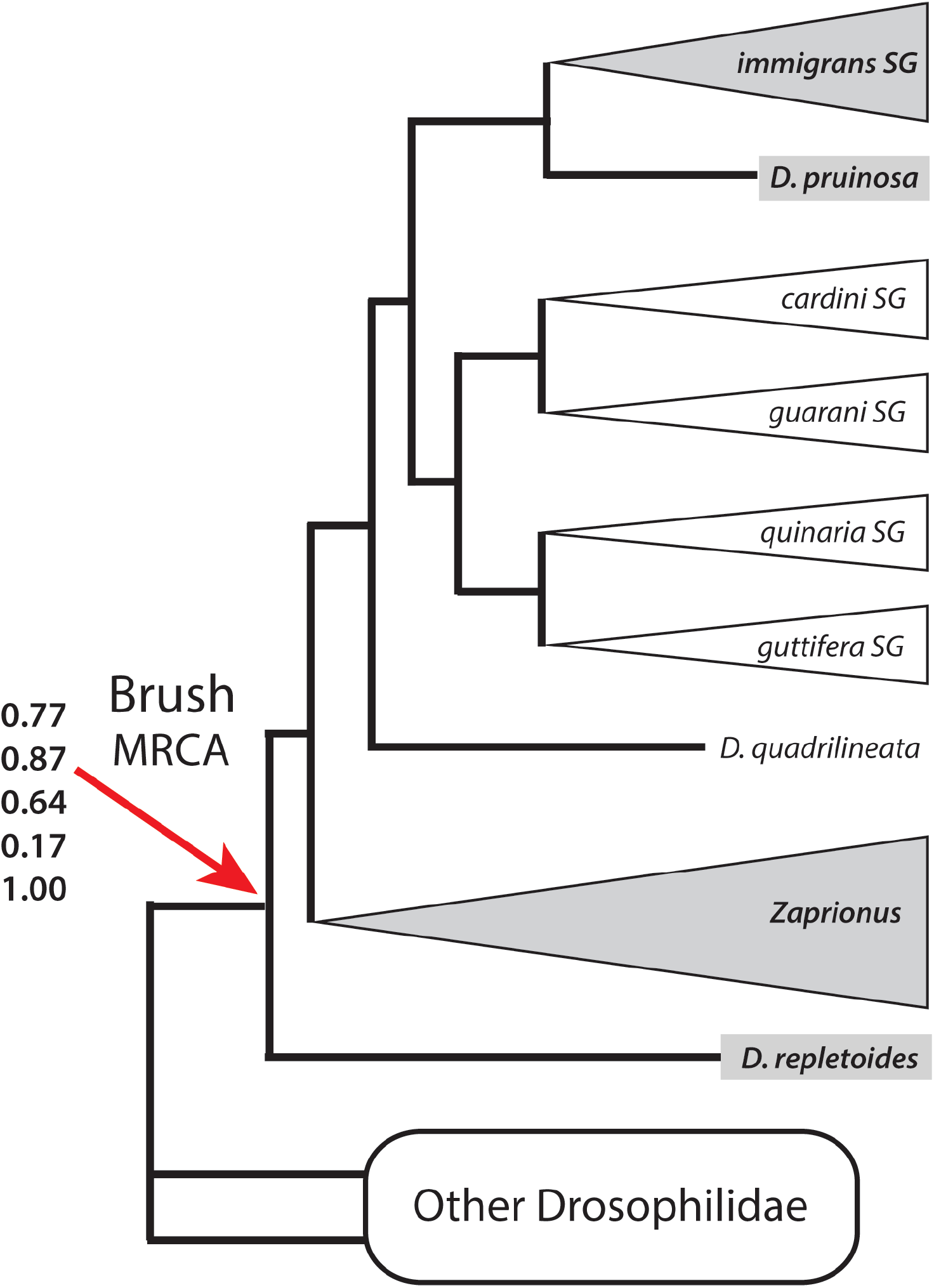
BUSCO phylogeny provides modest support for a single origin of the sex brush. This summary tree shows phylogenetic relationships among the four lineages that have the sex brush (grey background) and their brush-less relatives (white background). See Figure S3 for species-level phylogenetic relationships. Numbers at the “Brush MRCA” node denote the estimated probabilities that the last common ancestor of that clade had a sex brush, under five different models of trait evolution in the following order from top to bottom: MK model with unequal rates and strict molecular clock (see Fig. S5A for detailed reconstruction), MK model with unequal rates and random local relaxed clock (Fig. S5B), hidden states variable rates model with two latent rate classes (Fig. S5C), modified threshold model with 9 ordered latent states (Fig. S5D), and an approximation of Dollo model (Fig. S5E). See Methods and Table S4 for model descriptions.

In the end, the results of our phylogenetic analysis are inconclusive. While a single origin of the sex brush appears more likely than multiple independent gains based on the available species sample, the statistical support for this conclusion is model-dependent and not particularly strong, and it is possible that more comprehensive taxon sampling will produce a different result.

### The same cellular processes underlie sex brush development in all species

In *D. melanogaster*, TBR bristle precursors are specified between 6 and 12 hours after pupariation (AP) (Joshi and Orenic 2006, Schroff 2007). Initially, these cells are specified in sparse, loosely organized rows, and separated from one another by several epithelial cells. By 20-21 hrs AP, the bristle cells that are destined to make each TBR migrate toward each other to form a straight, contiguous row, while the intervening epithelial cells are expelled distally and proximally from the TBRs (Atallah et al. 2009; Tanaka et al. 2009). Previous work has shown that this developmental process can be modified in different ways to generate similar male-specific structures. For example, many species in the subgenera *Sophophora* and *Lordiphosa* bear sex combs arranged into a single longitudinal row along the proximo-distal leg axis. However, in some species the bristles that make up these sex combs are specified in longitudinal rows, at or near their final adult positions, while in others they develop originally as multiple TBRs which then rotate, align, and merge to form a single longitudinal row (Atallah et al. 2009; Tanaka et al. 2009; Atallah et al 2012).

In principle, a tightly packed brush in the ancestral TBR region could also develop and evolve through different mechanisms. For example, changes in cell fate specification could produce a higher bristle to epithelial cell ratio, leading to minimal spacing between the hair progenitor cells. Alternatively, the bristle cells and/or the intervening epithelial cells could migrate to increase the density of bristles in the brush area after cell fate specification is complete. Furthermore, a decrease in the size (or apical surface area) of epithelial cells in the brush could also lead to tighter bristle packing. The sex brushes of different species could potentially utilize different cellular mechanisms to produce similar adult structures.

To determine how sex brush development differs from that of the TBRs, we studied the species representing our four focal lineages: *D. pruinosa, D. immigrans, D. repletoides* and *Z. tuberculatus*. We used antibodies against membrane-localized proteins to visualize the shapes of bristle precursor cells and the surrounding epithelial cells during pupal leg development. When labeled with antibodies against the beta-catenin Armadillo (Arm) or the E-cadherin Shotgun (DE-cad), bristle cells can be distinguished from other cells by their unique membrane shape (Figure 4). We examined brush development at two timepoints: an early stage roughly corresponding to ~16-21 hr AP in *D. melanogaster*, when the bristle cells of the future TBRs begin to migrate toward each other and expel the intervening epithelial cells, and a later stage when cell migration is completed. We found that at the early stage, the brush region in all species already contains drastically higher numbers of bristle cells in comparison to the TBR region (e.g. see *Z. tuberculatus* female, Figure S5). Most brush bristles at this stage are each surrounded by four to six epithelial cells in all species. The neighboring bristle cells are separated from one another by one to two epithelial cells (Figure 4). At the late stage, this spacing appears to remain virtually invariant, although the cells become more regularly organized compared to the early stage (Figure 4). This lack of obvious cell migration is in contrast to the developing TBRs in the proximal region of the same segment (Figure 4) and in the homologous region in the female leg (e.g. *Z. tuberculatus*, Supplemental Figure S5).

**Figure 4.**
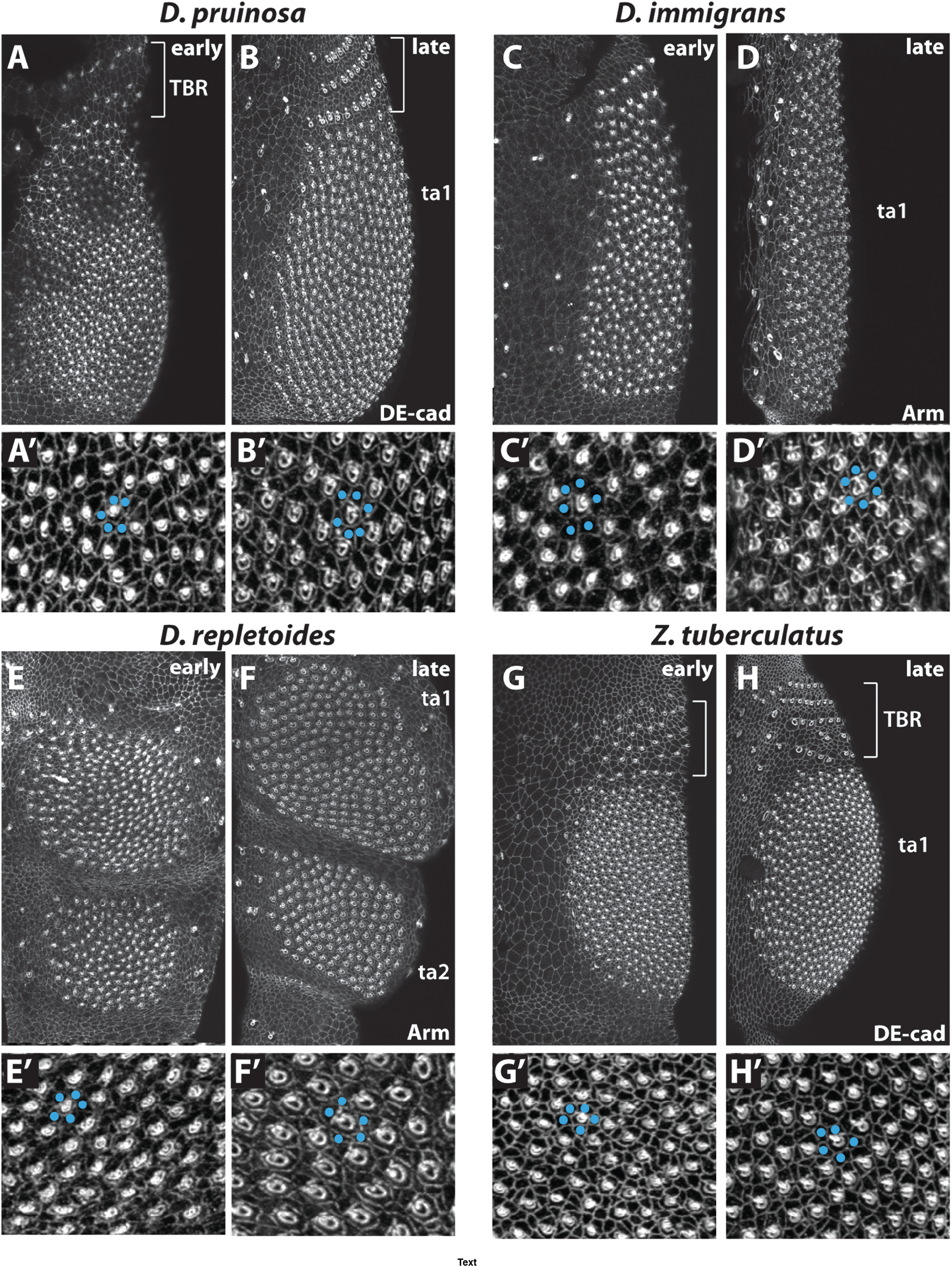
Sex brush development shows strong similarities across species. Developing brush hair cells were visualized by immunostaining for membrane markers E-cadherin (DE-cad) or Armadillo (Arm). For each species, the upper panels show the confocal projections of ta1 segment (A-D, G-H) or both ta1 and ta2 (E-F). The bottom panels (A’-H’) show close-up views. For each species, early developmental stages are shown on the left and later stages on the right. A-B’) *D. pruinosa*, 27 and 42 hrs after pupariation (AP). C-D’) *D. immigrans*, 24 and 43 hrs AP. E-F’) *D. repletoides*, 28 and 43 hrs AP. G-H’) *Z. tuberculatus*, 34 and 48 hrs AP. Hair progenitor cells can be distinguished from epithelial cells by a ring of bright staining with a dense punctum in the middle. In all four species, each hair cell is surrounded by 4-6 epithelial cells (blue dots), so that neighboring hair cells are separated from each other by 1-2 cells, at both the early and the late stages. The proximal TBRs (square brackets) form by expelling the intervening epithelial cells, with the bristle progenitor cells migrating closer together to make straight rows (Tanaka et al 2009) (e.g., compare A vs B, and G vs H). In contrast, no cell rearrangement is observed in the brush. For the early and late stages of each species, the numbers of legs analyzed were: *D. pruinosa* (5, 5), *Z. tuberculatus* (14, 11), *D. immigrans* (14, 10), *D. repletoides* (7, 6).

To determine if there are any changes in the ratio between bristle precursor and epithelial cells during brush development, we counted cells of each type in *Z. tuberculatus* and *D. immigrans* (Figure 5). We saw no significant change in this ratio between the early and late stages in *Z. tuberculatus* (epithelial/bristle cell ratio = 3.76 vs 3.73; Mann-Whitney U Test W=14.5, p-value = 0.7526). In *D. immigrans*, there was a slight but statistically significant *decrease* in the proportion of bristle cells (epithelial/bristle ratio = 3.21 vs 3.44; Mann-Whitney U Test W=0, p-value = 0.01167). Thus, there is no evidence for large-scale migration of bristle or epithelial cells that would bring bristle progenitors closer together as development proceeds.

**Figure 5.**
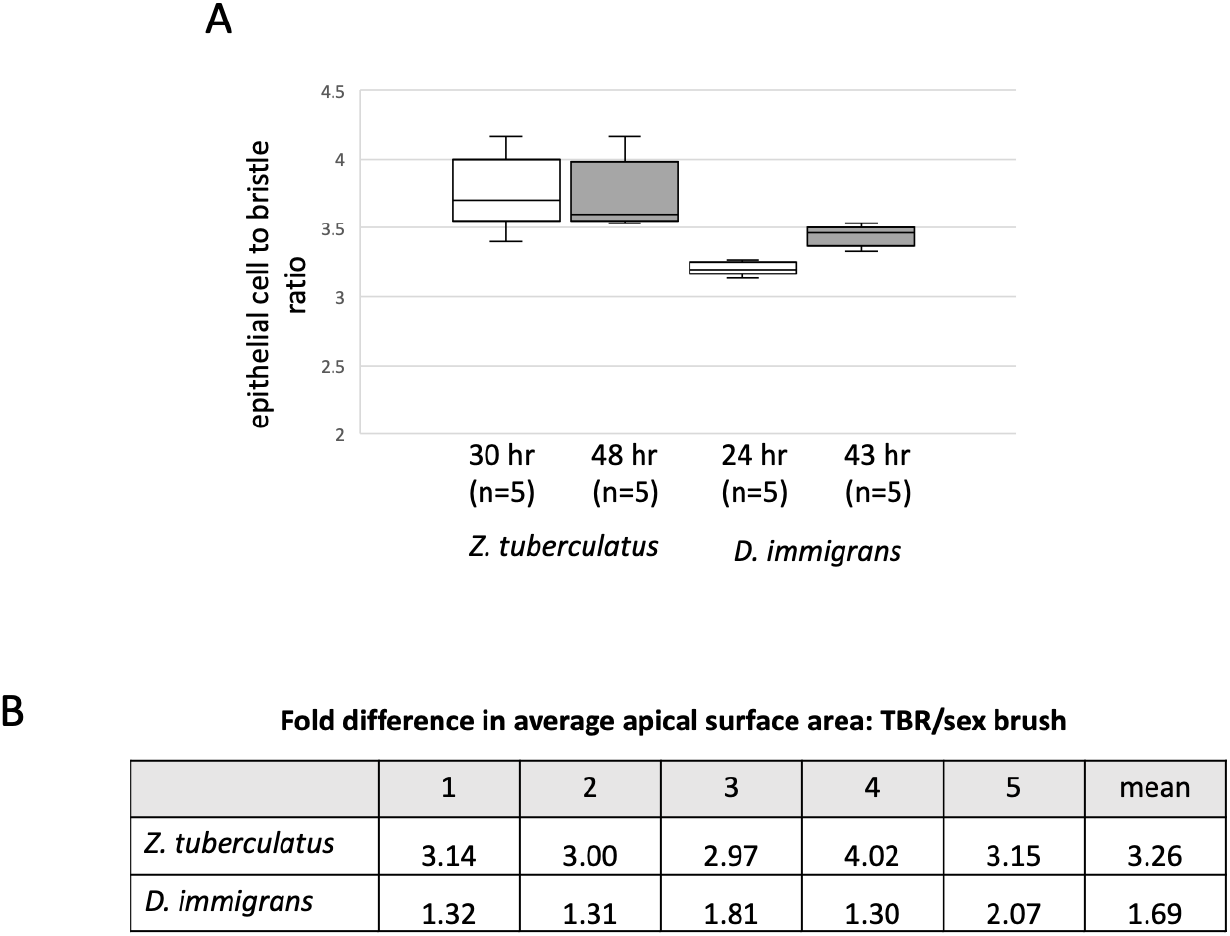
Analysis of cell density and cell size in the developing brush. A) Bristle density in the sex brush at early and late developmental stages in *Z. tuberculatus* and *D. immigrans*. The ratio between the number of epithelial cells and bristle cells was used as a proxy for bristle density. A group consisting of 30 bristle cells was selected from the brush area of each leg, and associated epithelial cells were counted. The absolute age (indicated in hours after pupariation) varies across species due to differences in the duration of pupal development. In *Z. tuberculatus*, the ratio of epithelial to bristle remained virtually unchanged, whereas, in *D. immigrans*, there was a slight increase (indicating a reduction in bristle density). B) Difference in epithelial cell size between the TBR region and the sex brush. Apical surface areas of 30 cells in each region were measured for five legs per species. The fold differences in the average size of epithelial cells in the TBR vs the brush regions are shown for individual legs.

The smaller cell size in the brush area may also contribute to the high density of hairs. Cell size difference between the brush area and the proximal TBR-bearing region of the same segment is particularly prominent in *Z. tuberculatus*. To quantitatively compare the size differences, we measured the apical surface area of 30 epithelial cells in each region of the tarsal segment in *Z. tuberculatus* and *D. immigrans* (n=5 legs per species). Although there were differences in the average apical surface area of cells among individual legs within each species, cells in the TBR regions were consistently larger than those in the sex brush (Figure 5; nested ANOVA, p-value <0.001 for both species). The average apical cell surface area in the TBR region was roughly three to four times larger than in the brush region for *Z. tuberculatus*, whereas in *D. immigrans* the fold difference was between 1.3 to 2.1 (Figure 5). In conclusion, the dense packing of hairs in the sex brush results both from the specification of bristle precursor cells at a high density and from a reduced cell size in the brush region, with a little to no contribution from processes such as cell migration and cell death.

## Discussion

The smallest clade to include all four brush-bearing lineages encompasses well over 1000 species (O’Grady and DeSalle, 2018; Kim et al, 2021), the vast majority of which lack sex brushes. Is this a result of a single origin followed by numerous losses, or is convergent evolution a more likely explanation? Unfortunately, even the largest phylogenetic dataset assembled to date for Drosophilidae does not provide a definitive answer to this question. Although a single origin appears more likely based on the available taxon sample, the support for this scenario is not very strong and depends on the choice of model of character evolution. Moreover, it is possible that our analyses underestimate the probability of multiple independent origins because the BUSCO dataset, although assembled without regard to leg morphology, substantially over-samples the brush-bearing species compared to their brush-less relatives.

The cellular mechanisms that produce sex brushes in different species appear essentially identical. In all species examined, the bristles that make up the sex brush are specified with only one or two intervening epithelial cells between them. Bristle specification in *Drosophila* and other insects is governed by a lateral inhibition mechanism, which is based on contact signaling between adjacent cells, and prevents two adjacent cells from both assuming the fate of bristle precursors (Simpson, 1990). Later in development, a tighter packing of bristles can be achieved through cell migration, as observed for example in the TBRs of the front legs (Atallah et al., 2009; Tanaka et al., 2009). In the case of sex brush, the future hairs appear to be specified at the maximum density allowed by lateral inhibition and involve little or no cell migration to produce the tight packing. Furthermore, in both species analyzed, the density of hairs is also increased by reduced cell size in the sex brush. Although the similarity of developmental mechanisms across species is consistent with a single origin of the sex brush, it cannot be seen as a clinching argument against convergent evolution. For example, similar cellular mechanisms of sex comb formation have evolved independently in distantly related lineages (Tanaka et al., 2009; Atallah et al., 2012).

We can only speculate about the selective forces that may drive the origin(s) and loss(es) of the sex brush. Males grasp female abdomens with their front legs in many *Drosophila* species, including those that lack any male-specific leg ornaments (Massey et al., 2019; Spieth, 1952). However, behavioral observations and functional experiments demonstrate that sex combs enhance grasping efficiency (Hurtado-Gonzales et al 2014; Massey et al. 2019; Spieth 1952). Our video recordings show that the proximal tarsal segments of the male T1 legs, including the sex brushes, are used to grab the female abdomen and resist the female’s efforts to dislodge the male during copulation attempts in *Z. tuberculatus*, *D. immigrans*, and *D. repletoides* (Supplemental movies 1-3). In all these species, females appear to resist mating attempts quite vigorously, using side-to-side bucking and wing vibrations. Thus, male sex brushes, which consist of hundreds of thin hairs that are hooked at the tips and have a very large combined surface area, could have evolved to provide a more secure grip of the female abdomen, especially if stronger grip is needed to counteract the female attempts to dislodge the male. It is easy to imagine how this type of mechanical advantage could provide the selective pressure for the origin (and perhaps several convergent origins) of the sex brush.

If so, why don’t all *Drosophila* species evolve sex combs, brushes, or other grasping structures? Development does not provide any clues. At the level of cell biology, both the ancestral/female condition (cell migration that produces tightly packed bristle rows from sparsely spaced precursors) and the derived/male condition (specification of bristle precursors at the maximum density permitted by lateral inhibition) are the same in all lineages where the sex brush is present. There is no a priori reason to think that the transition between these modes of development is easier in some species than others. The answer may lie instead in either behavior or population genetics. Although males of different species use their sex brushes in similar ways, we don’t know the female side of the story. If females of different species vary in their responses to male grasping, the evolution of specialized leg structures in males may not be universally favored. This may also explain why both the sex brushes and the sex combs (Kopp, 2011) have been secondarily lost multiple times. Moreover, it is difficult to know whether the female preferences observed today are the same as they were in the distant past when the male-specific structures evolved (Watts et al., 2019). A systematic phylogenetic analysis of mating behavior, including a dense sampling of lineages that lack male-specific leg modifications, will be needed to test whether morphological evolution correlates with the evolution of behavior.

Alternatively, the origin of a new trait such as the leg brush may require such an unlikely series of genetic changes that it may often fail to occur even in response to strong selective pressure. For example, it is possible that while a single mutation is sufficient to modify or eliminate an existing morphological structure, the origin of a *new* structure may require simultaneous changes in multiple genes. From the population-genetic perspective, this would mean that functionally novel and positively interacting alleles at multiple loci must segregate in the same population at the same time in order for selection in favor of a new structure to be effective. Naturally, this would greatly reduce the probability of evolutionary innovations compared to other types of phenotypic change. We hope that research models where both the functional roles and the genetic basis of novel traits can be studied in parallel will help elucidate why certain innovations appear in some lineages but fail to evolve in others.

## Supporting information

Supplemental Figures and Tables except Table S3

## Declarations

### Funding

This work was supported by NIH grant 5R35GM122592 to AK, NIH grant F32 GM135998 to BYK, and by funds from Ludwig-Maximilians-Universität München to Nicolas Gompel, in whose lab some of the work was conducted. The Olympus FV1000 confocal used in this study was purchased using NIH Shared Instrumentation Grant 1S10RR019266-01.

### Competing interests

The authors declare that they have no competing interests.

### Availability of data and material

The datasets used in this study are included in the manuscript or will be made available in GenBank upon publication.

### Code availability

Not applicable

### Author’s contributions

AK conceived the study. KT, OB, BYK, AK and JHM performed experiments and collected data. KT, BYK, AS, AT, and AK analyzed the data. KT and AK wrote the manuscript with contributions from AT, BYK, and AS.

### Ethics approval

Not applicable

### Consent to participate

Not applicable

### Consent for publication

Not applicable

## Acknowledgements

We thank the US *Drosophila* species stock center, Ehime University *Drosophila* stock center, J.-R. David, S. Prigent, and M. Watada for *Drosophila* strains and specimens; Developmental Studies Hybridoma Bank for antibodies; A. Comeault, A. Gloss, M. Lang, D. Matute, D. Miller, V. Orgogozo, and N. Whiteman for genome assemblies; M. Toda for advice on *Zaprionus* phylogeny; G. Rice for *D. immigrans* images; and A. Yassin for comments on the manuscript. We also thank the MCB Light Microscopy Imaging Facility, Department of Cell Biology and Human Anatomy EM Lab, and Materials Science and Engineering AMCaT Lab at UC Davis for equipment and imaging assistance, and Gwyneth Card and W. Ryan Williamson for use of the Photron camera. AK and OB are grateful to N. Gompel for allowing some of this work to be conducted in his lab while on a sabbatical visit, and to members of the Gompel lab for treating them as their labmates.

## Notes

### Competing Interest Statement

The authors have declared no competing interest.

### Summary of Updates

1) The current version of the manuscript contains additional phylogenetic analyses for which we assembled considerably larger dataset (250 genes from 207 Drosophila species). The results of these analyses are less definitive about the evolutionary origins of our focal trait than the smaller phylogenetic study in the previous version of the manuscript. Whereas the earlier analysis supported the convergent origin, the new analyses favor a single origin followed by multiple losses of that trait; however, support for this scenario is not very strong, and convergent evolution cannot be ruled out with any confidence either. We have revised the results and discussion sections of the manuscript accordingly. 2) The title and the abstract have been changed to reflect new conclusions. 3) The manuscript has three additional authors. 4) The result section contains the phylogenetic analyses with the larger dataset as well as the ancestral state reconstruction. 5) We also conducted more thorough quantitative analyses of cell dynamics during development. 6) The previous version of the manuscript should not be cited since its conclusions are superseded by the new analyses in the current version.

