## Supplemental Information for "Evolution and development of male-specific leg brushes in Drosophilidae"

### Supplemental text – notes on Drosophila taxonomy

The *immigrans* species group

The *immigrans* species group and the wider *immigrans-tripunctata* radiation have long played a pivotal role in *Drosophila* systematics (Throckmorton, 1975; Yassin, 2013). However, the composition of the *immigrans* group has not been entirely clear. Historically, this group was proposed to include five subgroups: *immigrans, hypocausta, nasuta, quadrilineata*, and *curviceps* (Huang et al., 2002; Zhang and Toda, 1988, 1992; Zhang et al., 1995). Recent studies have shown, however, that both the *hypocausta* and the *immigrans* subgroups are likely polyphyletic, as some species assigned to these subgroups are actually closer to the *nasuta* subgroup (Da Lage et al., 2007; Katoh et al., 2007; Rice et al., 2018). The *quadrilineata* and *curviceps* subgroups have been the source of even greater complications. Previous phylogenetic studies have provided evidence against a sister-group relationship between *D. quadrilineata* and the rest of the *immigrans* species group (Katoh et al., 2007; Morales-Hojas and Vieira, 2012; Yassin, 2013). Moreover, *D. annulipes*, traditionally assigned to the *quadrilineata* subgroup (Lin and Tseng, 1973), was found to be distantly related to *D. quadrilineata*, and closer to the *virilis-repleta* radiation (Katoh et al., 2007). The phylogeny of Huang et al (Huang et al., 2002) showed a close relationship between *D. annulipes* and the clade composed of *D. curviceps* and *D. oritisa*, another species assigned to the *curviceps* subgroup. Morales-Hojas and Vieira (Morales-Hojas and Vieira, 2012) confirmed the distant relationship between *D. annulipes* and *D. quadrilineata*, as well as between *D. annulipes* and the *immigrans* species group *sensu stricto*. Yassin (Yassin, 2013) pointed out important differences between *D. quadrilineata* and the *curviceps* subgroup and the *immigrans* group *s.s*. Finally, Pradhan et al (Pradhan et al., 2015) removed the *curviceps* subgroup from the *immigrans* species group, elevating it to the status of a separate species group.

Our analysis, based on a larger amount of sequence data than previous studies, confirms these observations, namely that (1) *D. quadrilineata* is not closely related either to the *immigrans* species group *s.s.* or to *D. annulipes*, (2) *D. curviceps* is not closely related to the *immigrans* species group *s.s.*, and (3) *D. annulipes* is related to *D. curviceps*. Based on these results, we support the conclusions of other authors (Katoh et al., 2007; Pradhan et al., 2015; Yassin, 2013) that the definition of the *immigrans* species group should be restricted to the *immigrans*, *hypocausta*, and *nasuta* subgroups. Further work, with a more extensive sampling of the *immigrans* group *s.s.*, is needed to revise its internal taxonomy and re-establish monophyletic subgroups. Similarly, better taxon sampling will be needed to determine whether *D. annulipes* should be reassigned to the *curviceps* species group (Pradhan et al., 2015), or whether it is possible to establish a monophyletic *quadrilineata* species group.

*Zaprionus* and *Anaprionus*

In *Zaprionus*, male brushes are present in most African species, with the exception of *Z. neglectus*, *Z. spineus*, and *Z. spinosus* (Tsacas and Chassagnard, 1990; Yassin et al., 2008; Yassin and David, 2010). The *Zaprionus* phylogeny is not fully resolved, but the distant relationship between the first species and the last two suggests that their lack of brushes is likely to reflect independent secondary losses. The situation is more complicated among species assigned to the Oriental *Anaprionus* subgenus of *Zaprionus*. Many of its members, including *Z. lineosus, Z. spinilineosus, Z. orissaensis, Z. multistriatus, Z. grandis*, and *Z. aungsani*, lack leg brushes (Gupta, 1972; Kikkawa and Peng, 1938; Okada and Carson, 1983; Wynn and Toda, 1988). However, *Anaprionus* is now thought to be polyphyletic (Yassin, 2007; Yassin et al., 2010), and these species appear to be more closely related to the genus *Xenophorticella* than to *Zaprionus* sensu stricto (M. Toda, pers. comm.). Other *Anaprionus* species such as *Z. bogoriensis, Z. obscuricornis,* and *Z. pyinoolwinensis* have leg brushes (Mainx, 1958; Okada, 1964; Wynn and Toda, 1988) and likely form a clade with the Afrotropical *Zaprionus* (M. Toda, pers. comm.). Our analysis confirms that the Asian *Z.* (*Anaprionus*) *bogoriensis* is the closest outgroup to the African *Zaprionus* species (Figure S3). Future phylogenetic analyses with comprehensive sampling of Oriental *Anaprionus* species should refine our understanding of sex brush evolution in *Zaprionus*.

*loiciana* species complex

The *loiciana* species complex, which is not currently assigned to any species group within *Drosophila*, contains six described species: *D. pruinosa, D. loiciana, D. pachneissa, D. semipruinosa, D. allochroa, and D. xanthochroa* (Tsacas, 2002; Tsacas and Chassagnard, 2000). All these species have male sex brushes of different sizes.

*D. repletoides*

*D. repletoides* does not have any known close relatives. It is assigned to the *tumiditarsus* species group (O'Grady and DeSalle 2018); however, the species described originally as *D. tumiditarsus* (Tan et al., 1949) was later synonymized with *D. repletoides* (Hsu, 1943; Wheeler, 1981). Yassin (Yassin, 2007) suggested that some species currently classified as *Zaprionus* (*Z. multistriatus*, *Z. flavofasciatus*, and *Z. cercociliaris*) could in fact be more closely related to *D. repletoides* than to *Zaprionus*. Unfortunately, these species have not been included in any molecular phylogenies, and we lack appropriate specimens.

### Supplemental Figures

**Supplemental Figure S1.** Strict consensus of 11 trees with the cumulative posterior probability of 95% for the 8-locus dataset, labeled as in Figure 2. Numbers at each node indicate the posterior probabilities of the respective taxon bipartitions.

**Supplemental Figure S2.** Bayesian phylogenetic tree for the 8-locus dataset, with the dataset partitioned by gene and codon.

**Supplemental Figure S3.** A maximum likelihood tree of 207 drosophilid species reconstructed using IQ-TREE v1.6.5 from 250 single-copy BUSCO loci (572,343 total sites) (Table S3) and rooted with *Anopheles gambiae*. At each node, the three measures of node support are, in order: 1000 replicates of ultrafast bootstrap (Minh et al., 2013), a Bayesian-like transformation of approximate likelihood ratio test (aLRT) (Anisimova et al., 2011), and aLRT with the nonparametric Shimodaira–Hasegawa correction (SH-aLRT). The *A. gambiae* branch is not to scale. See Methods for details of phylogeny reconstruction.

**Supplemental Figure S4.** Ancestral state reconstruction under five different models of trait evolution. The phylogenetic tree is based on 250 single-copy BUSCO genes extracted from whole genome sequences (Supplemental Table S3). Species are colored according to sex brush state (red = present, blue = absent). Pie charts at internal nodes reflect estimated probabilities of ancestral character states under five different models of trait evolution: (A) MK model with unequal rates and a strict molecular clock (Lewis, 2001); (B) MK model with unequal rates and a random local relaxed clock (Drummond and Suchard, 2010); (C) Hidden states variable rates model with two latent rate classes (Beaulieu et al., 2013); (D) Modified threshold model (Felsenstein, 2005) with 9 ordered latent states; (E) Approximation of a Dollo model, with rate of loss >300-fold higher than rate of gain. See Methods and Table S4 for details of ancestral character reconstruction.

**Supplemental Figure S5.** TBR development in female *Z. tuberculatus*. Developing bristle and epithelial cells were visualized by immunostaining for E-cadherin (DE-cad) at an early stage (30 hr AP, left) and a later stage (48 hr AP, right). Bristle progenitor cells can be distinguished from epithelial cells by a ring of bright staining with a dense punctum in the middle. The upper panels show the confocal projections of the ta1 segment (A, B). The bottom panels (C, D) show magnified views of the distal TBR region from the upper panels. As in the male legs, the TBRs form by expelling the intervening epithelial cells, with the bristle progenitor cells migrating closer together to make straight rows.

### Supplemental Tables

**Supplemental Table S1.** Sequence data used in the 8-locus phylogenetic analysis.

**Supplemental Table S2.** Primers used to amplify gDNA sequences for the 8-locus phylogenetic analysis.

**Supplemental Table S3.** Species and BUSCO loci for the 250-locus phylogenetic analysis.

**Supplemental Table S4.** Model parameters used in ancestral character reconstruction.

### Supplemental movies 1-3

Copulation behavior in *D. immigrans* (Movie 1)*, D. repletoides* (Movie 2), and *Z. tuberculatus* (Movie 3). Previous work in the *Drosophila melanogaster* species group has shown that males use their sex combs, which are also located on the T1 tarsus, to grab the female genitalia as in *D. melanogaster,* or the middle abdominal segments as in *D. kikkawai, D. ananassae*, and *D. bipectinata* (Massey et al., 2019). In these species, male T2 and T3 legs remain on the substrate and are not involved in mating. Typically, these males proceed very quickly from mounting to attempted copulation; females may resist by walking away or using their hind legs to kick the male off. In contrast, *D. immigrans, Z. tuberculatus*, and *D. repletoides* males use their T1 legs to grab females more anteriorly, near the constriction between the thorax and abdomen. In the former two species (Movies 1 and 2), males also use their T2 legs to grab the female mid-abdomen, while in *D. repletoides* T1, T2 and T3 legs are all used to grab the female so that the male "rides" on the female and is not in contact with the substrate (Movie 3). In all these species, females appear to resist mating attempts more vigorously than in the *melanogaster* group, using side-to-side bucking and wing vibrations in apparent efforts to dislodge the male, while the males use their legs to resist these efforts. The delay between mounting and attempted copulation is longer in the brush-bearing species, especially in *Z. tuberculatus*, than in the comb-bearing species; most mountings result in the male being eventually dislodged and do not lead to copulation attempts.
