## Supplemental Figures and Tables except Table S3 for "Evolution and development of male-specific leg brushes in Drosophilidae"

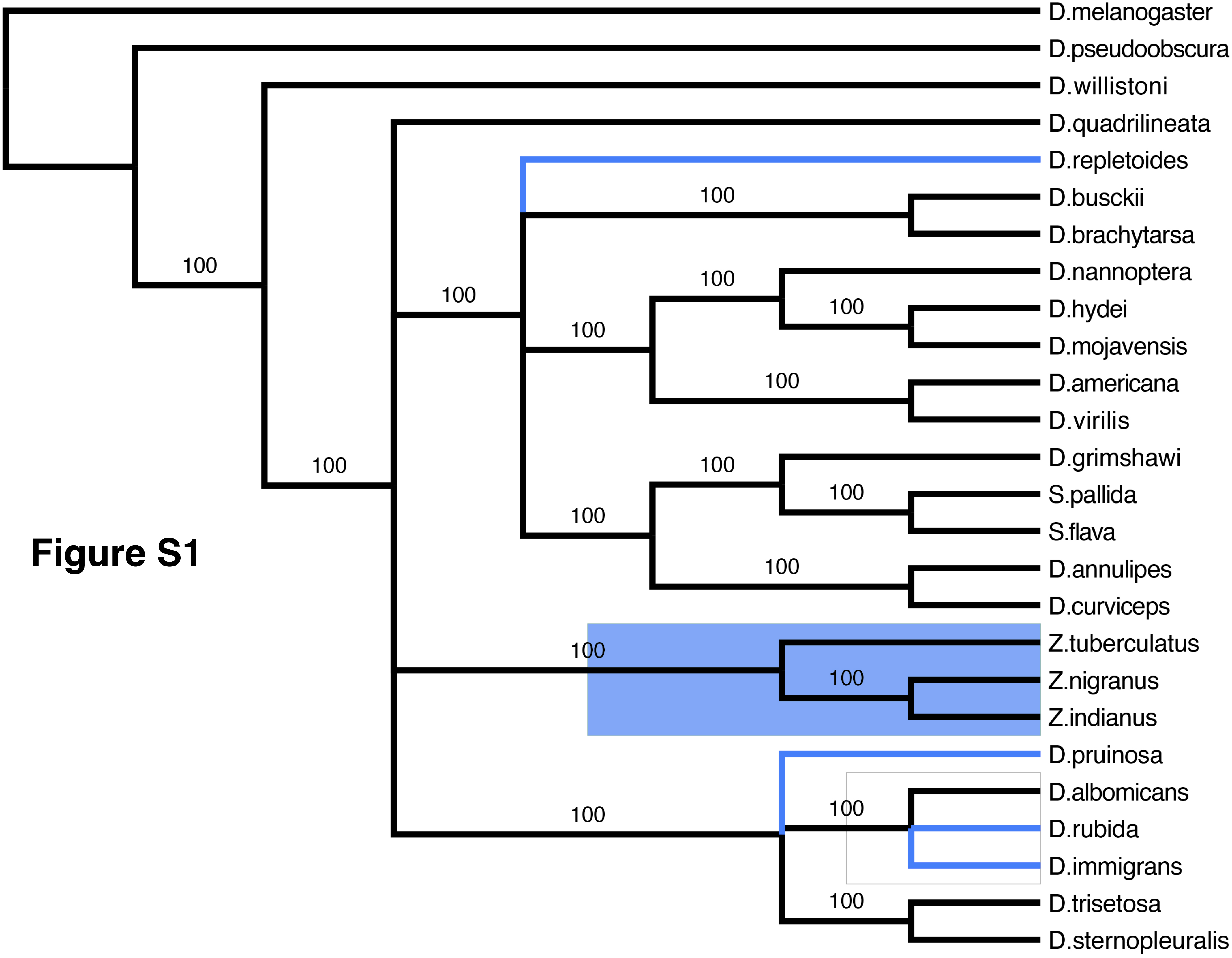
D.melanogaster
D.pseudoobscura
D.willistoni
D.quadrilineata
D.repletoides
D.busckii
D.brachytarsa
D.nannoptera
D.hydei
D.mojavensis
D.americana
D.virilis
D.grimshawi
S.pallida
S.flava
D.annulipes
D.curviceps
Z.tuberculatus
Z.nigranus
Z.indianus
D.pruinosa
D.albomicans
D.rubida
D.immigrans
D.trisetosa
D.sternopleuralis
100
100
100
100
100
100
100
100
100
100
100
100
100
100
100
100
100
100

**Figure S1**

**Figure S2**

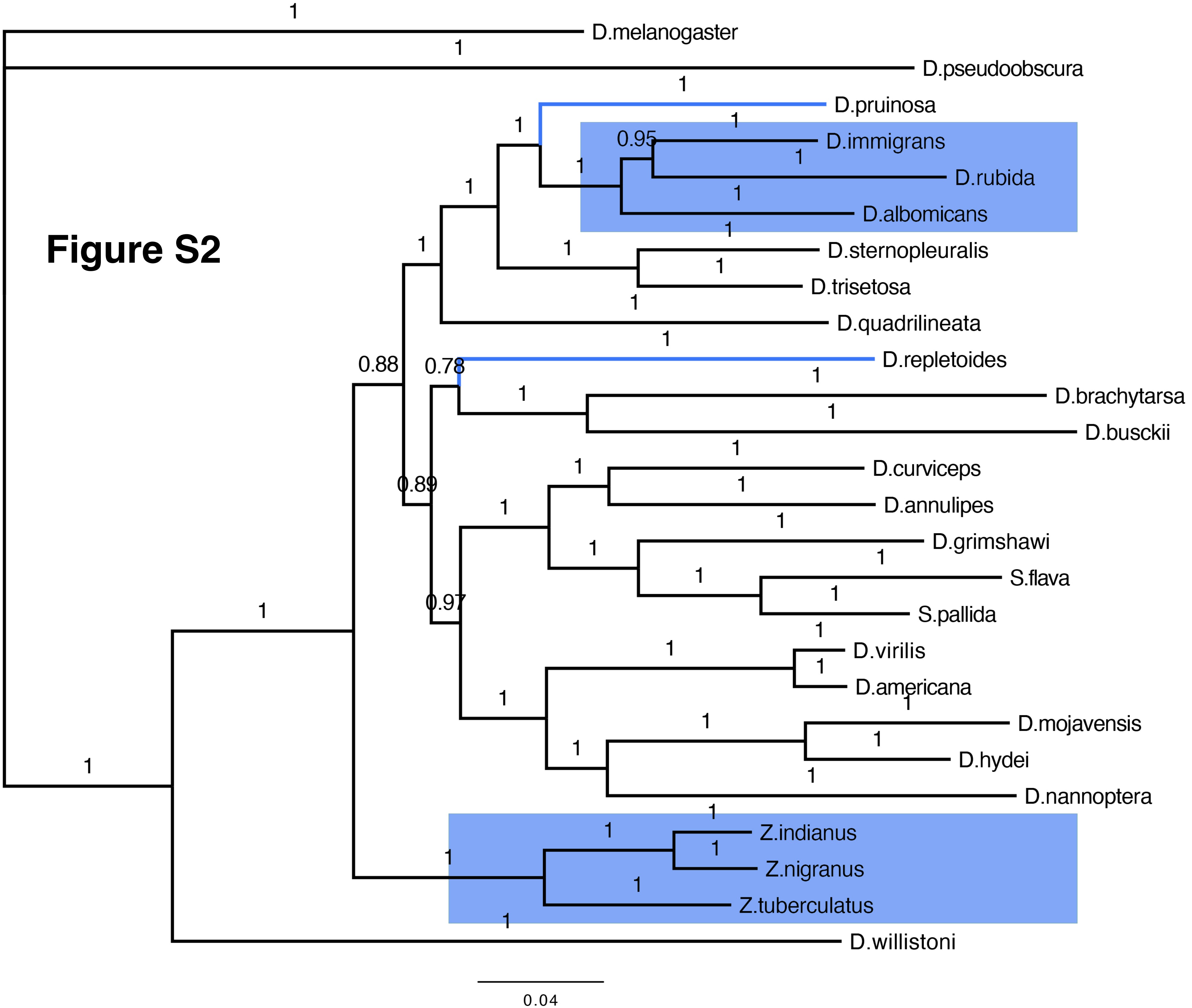
D.melanogaster
D.pseudoobscura
D.pruinosa
D.immigrans
D.rubida
D.albomicans
D.sternopleuralis
D.trisetosa
D.quadrilineata
D.repletoides
D.brachytarsa
D.busckii
D.curviceps
D.annulipes
D.grimshawi
S.flava
S.pallida
D.virilis
D.americana
D.mojavensis
D.hydei
D.nannoptera
Z.indianus
Z.nigranus
Z.tuberculatus
D.willistoni
0.04

**Figure S3**

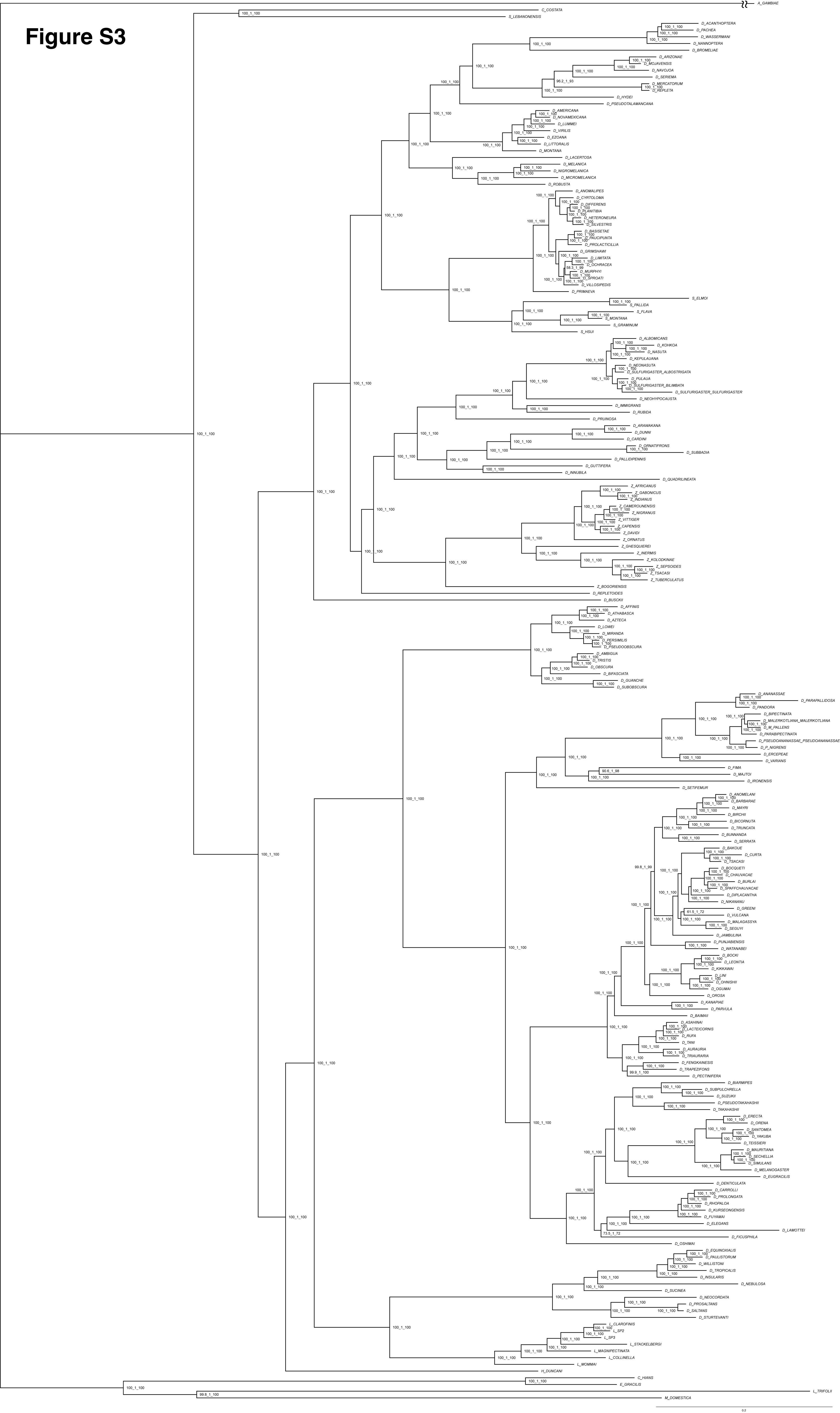
A_GAMBIAE
C_COSTATA
S_LEBANONENSIS
D_ACANTHOPTERA
D_PACHEA
D_WASSERMANI
D_NANNOPTERA
D_BROMELIAE
D_ARIZONAE
D_MOJAVENSIS
D_NAVOJOA
D_SERIEMA
D_MERCATORUM
D_REPLETA
D_HYDEI
D_PSEUDOTALAMANCANA
D_AMERICANA
D_NOVAMEXICANA
D_LUMMEI
D_VIRILIS
D_EZOANA
D_LITTORALIS
D_MONTANA
D_LACERTOSA
D_MELANICA
D_NIGROMELANICA
D_MICROMELANICA
D_ROBUSTA
D_ANOMALIPES
D_CYRTOLOMA
D_DIFFERENS
D_PLANITIBIA
D_HETERONEURA
D_SILVESTRIS
D_BASISETAE
D_PAUCIPUNTA
D_PROLACTICILLIA
D_GRIMSHAWI
D_LIMITATA
D_OCHRACEA
D_MURPHYI
D_SPROATI
D_VILLOSIPEDIS
D_PRIMAEVA
S_ELMOI
S_PALLIDA
S_FLAVA
S_MONTANA
S_GRAMINUM
S_HSUI
D_ALBOMICANS
D_KOHKOA
D_NASUTA
D_KEPULAUANA
D_NEONASUTA
D_SULFURIGASTER_ALBOSTRIGATA
D_PULAUA
D_SULFURIGASTER_BILIMBATA
D_SULFURIGASTER_SULFURIGASTER
D_NEOHYPOCAUSTA
D_IMMIGRANS
D_RUBIDA
D_PRUINOSA
D_ARAWAKANA
D_DUNNI
D_CARDINI
D_ORNATIFRONS
D_SUBBADIA
D_PALLIDIPENNIS
D_GUTTIFERA
D_INNUBILA
D_QUADRILINEATA
Z_AFRICANUS
Z_GABONICUS
Z_INDIANUS
Z_CAMEROUNENSIS
Z_NIGRANUS
Z_VITTIGER
Z_CAPENSIS
Z_DAVIDI
Z_ORNATUS
Z_GHESQUIEREI
Z_INERMIS
Z_KOLODKINAE
Z_SEPSOIDES
Z_TSACASI
Z_TUBERCULATUS
Z_BOGORIENSIS
D_REPLETOIDES
D_BUSCKII
D_AFFINIS
D_ATHABASCA
D_AZTECA
D_LOWEI
D_MIRANDA
D_PERSIMILIS
D_PSEUDOOBSCURA
D_AMBIGUA
D_TRISTIS
D_OBSCURA
D_BIFASCIATA
D_GUANCHE
D_SUBOBSCURA
D_ANANASSAE
D_PARAPALLIDOSA
D_PANDORA
D_BIPECTINATA
D_MALERKOTLIANA_MALERKOTLIANA
D_M_PALLENS
D_PARABIPECTINATA
D_PSEUDOANANASSAE_PSEUDOANANASSAE
D_P_NIGRENS
D_ERCEPEAE
D_VARIANS
D_FIMA
D_MAJTOI
D_IRONENSIS
D_SETIFEMUR
D_ANOMELANI
D_BARBARAE
D_MAYRI
D_BIRCHII
D_BICORNUTA
D_TRUNCATA
D_BUNNANDA
D_SERRATA
D_BAKOUE
D_CURTA
D_TSACASI
D_BOCQUETI
D_CHAUVACAE
D_BURLAI
D_SPAFFCHAUVACAE
D_DIPLACANTHA
D_NIKANANU
D_GREENI
D_VULCANA
D_MALAGASSYA
D_SEGUYI
D_JAMBULINA
D_PUNJABIENSIS
D_WATANABEI
D_BOCKI
D_LEONTIA
D_KIKKAWAI
D_LINI
D_OHNISHII
D_OGUMAI
D_OROSA
D_KANAPIAE
D_PARVULA
D_BAIMAII
D_ASAHINAI
D_LACTEICORNIS
D_RUFA
D_TANI
D_AURAURIA
D_TRIAURARIA
D_FENGKAINESIS
D_TRAPEZIFONS
D_PECTINIFERA
D_BIARMIPES
D_SUBPULCHRELLA
D_SUZUKII
D_PSEUDOTAKAHASHII
D_TAKAHASHII
D_ERECTA
D_ORENA
D_SANTOMEA
D_YAKUBA
D_TEISSIERI
D_MAURITIANA
D_SECHELLIA
D_SIMULANS
D_MELANOGASTER
D_EUGRACILIS
D_DENTICULATA
D_CARROLLI
D_PROLONGATA
D_RHOPALOA
D_KURSEONGENSIS
D_FUYAMAI
D_ELEGANS
D_LAMOTTEI
D_FICUSPHILA
D_OSHIMAI
D_EQUINOXIALIS
D_PAULISTORUM
D_WILLISTONI
D_TROPICALIS
D_INSULARIS
D_NEBULOSA
D_SUCINEA
D_NEOCORDATA
D_PROSALTANS
D_SALTANS
D_STURTEVANTI
L_CLAROFINIS
L_SP2
L_SP3
L_STACKELBERGI
L_MAGNIPECTINATA
L_COLLINELLA
L_MOMMAI
H_DUNCANI
C_HIANS
E_GRACILIS
L_TRIFOLII
M_DOMESTICA
0.2

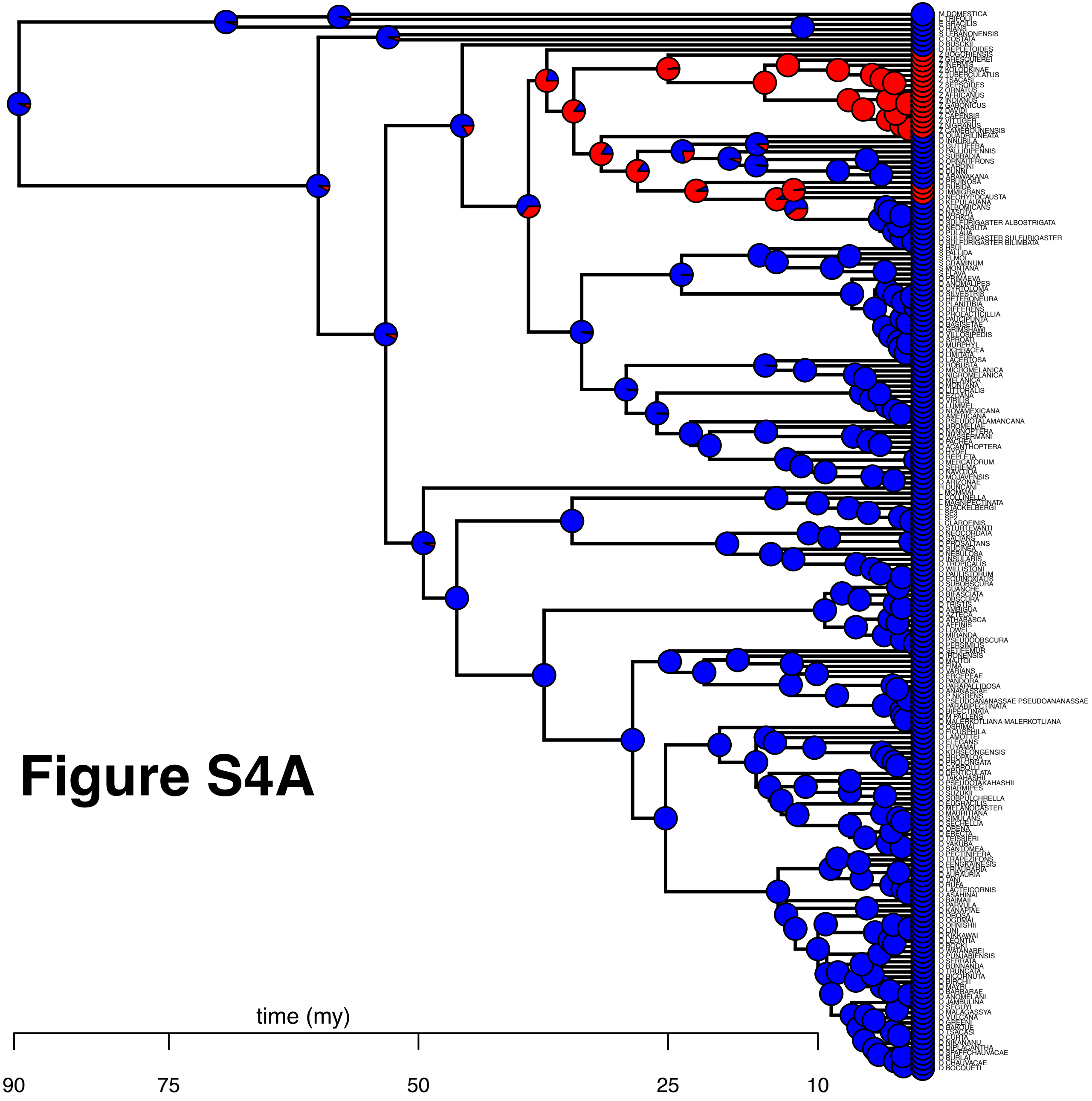

Figure S4A

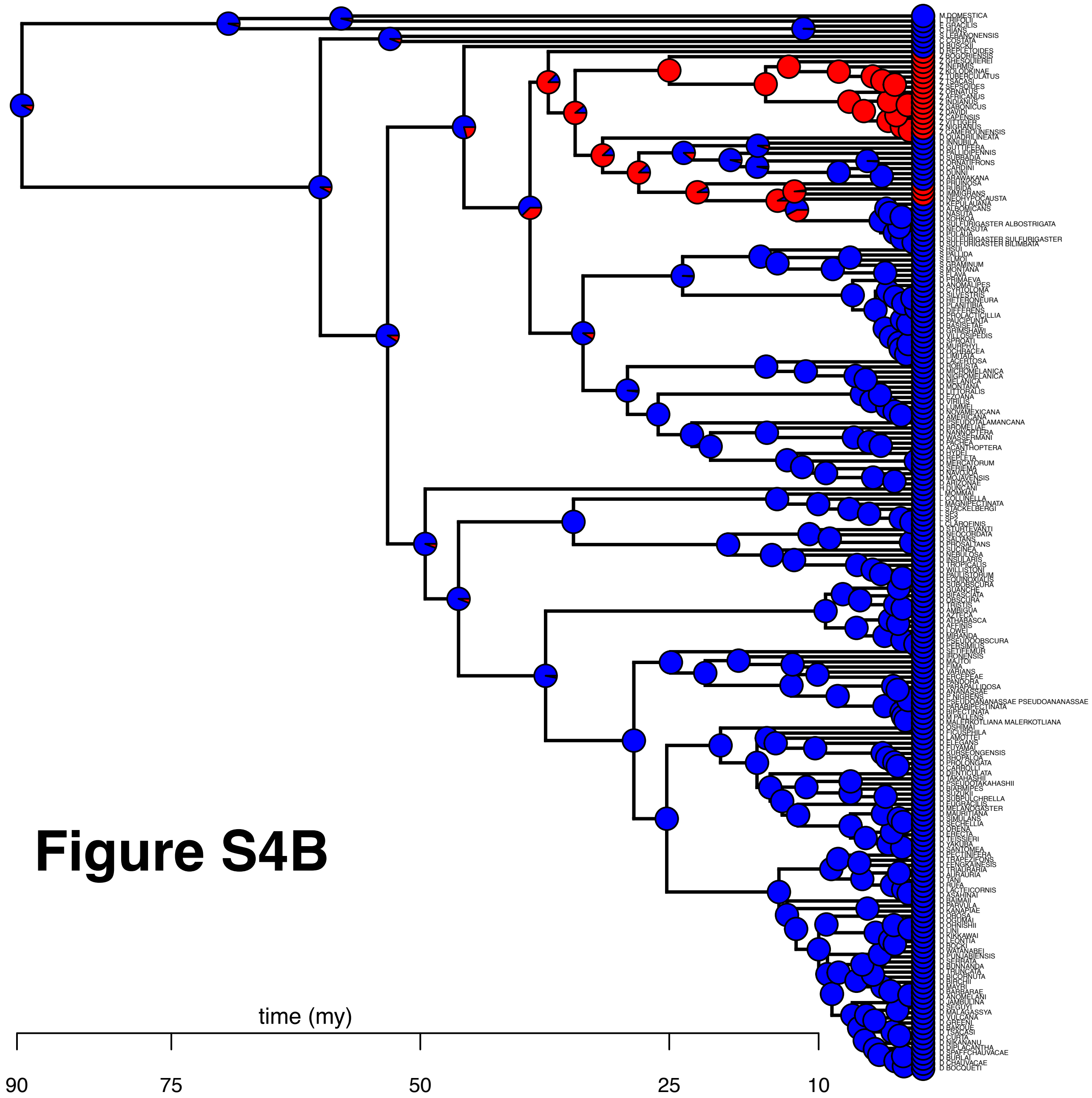

Figure S4B

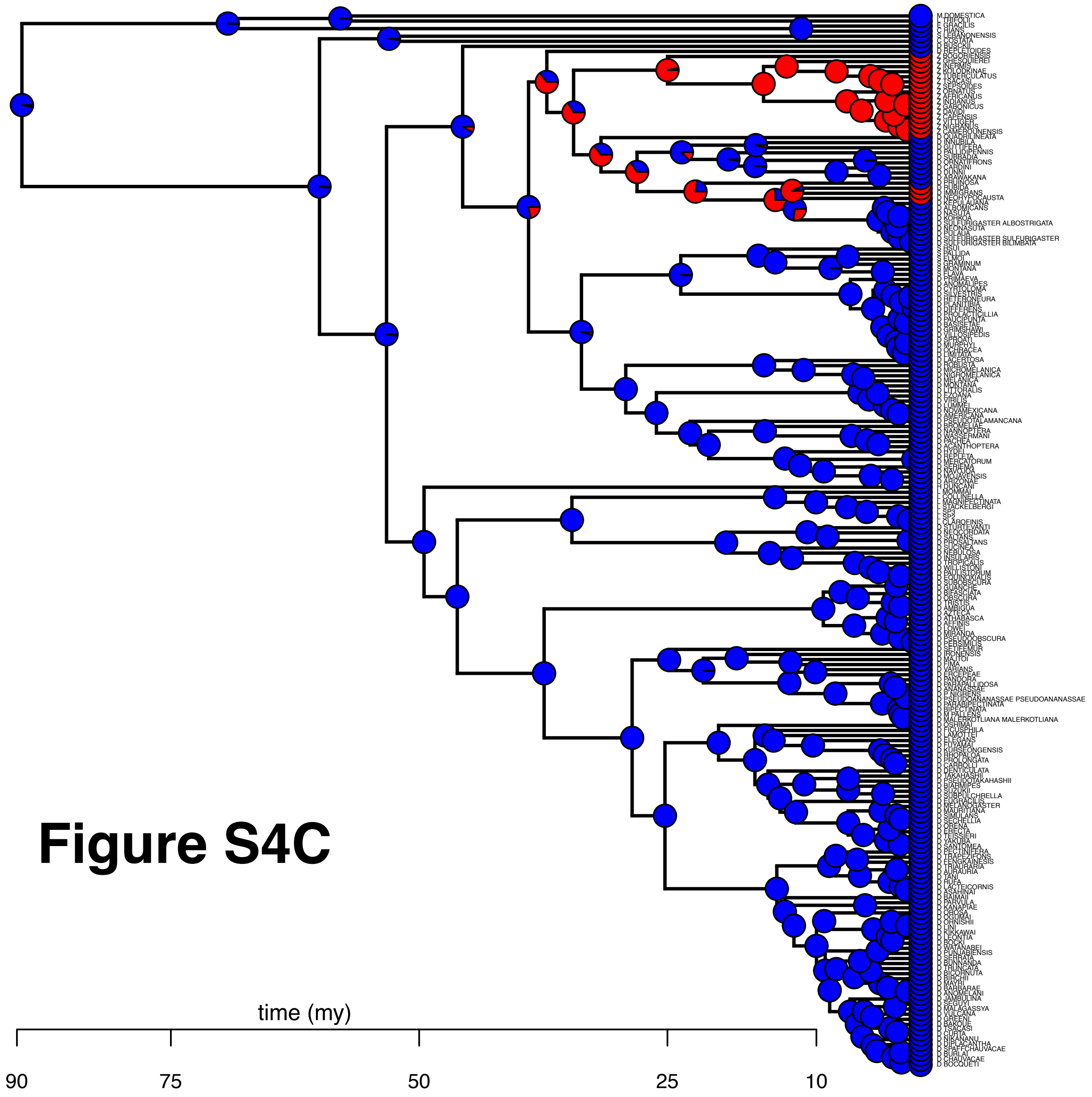

### Figure S4C

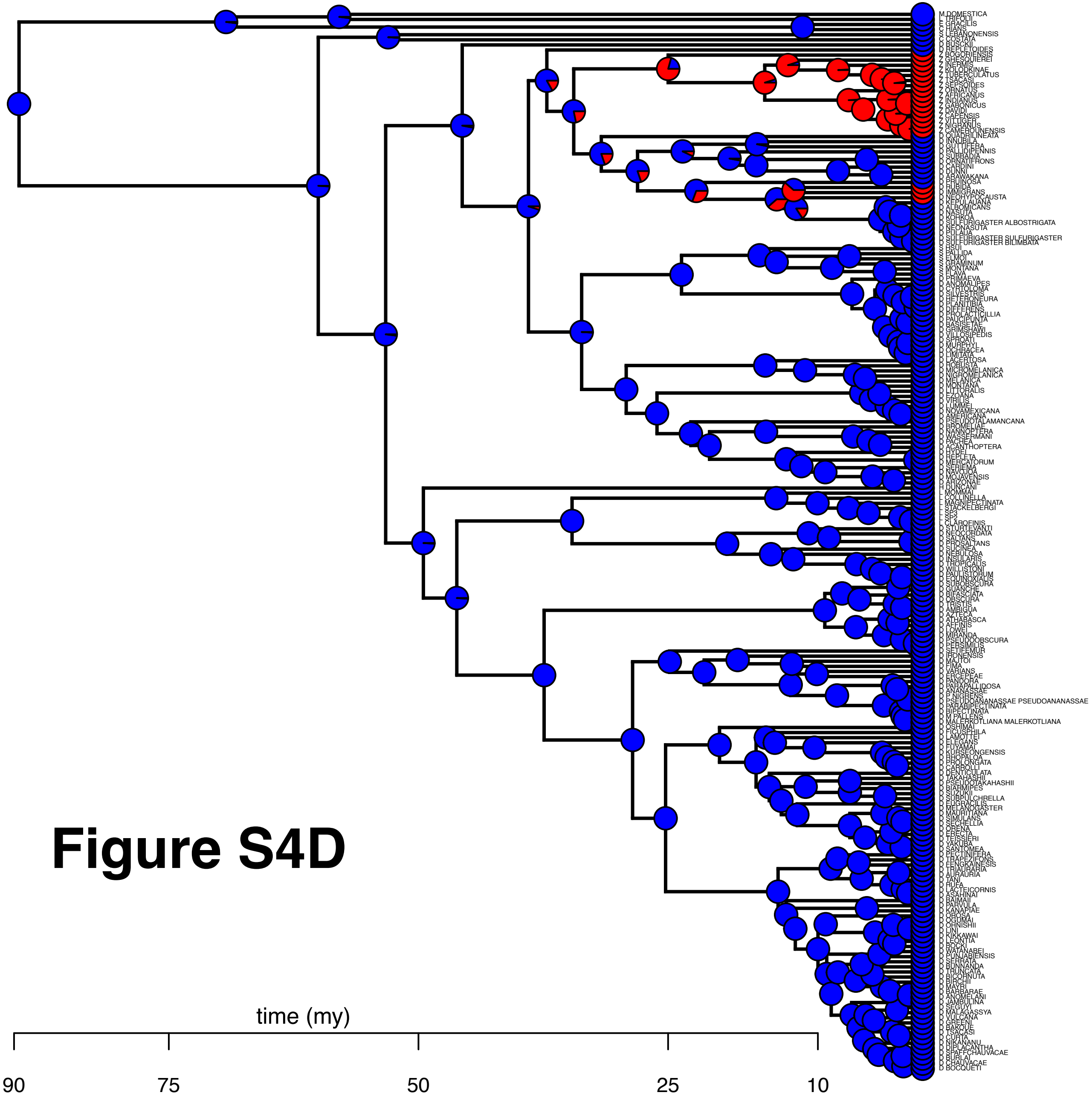

**Figure S4D**

### Figure S5

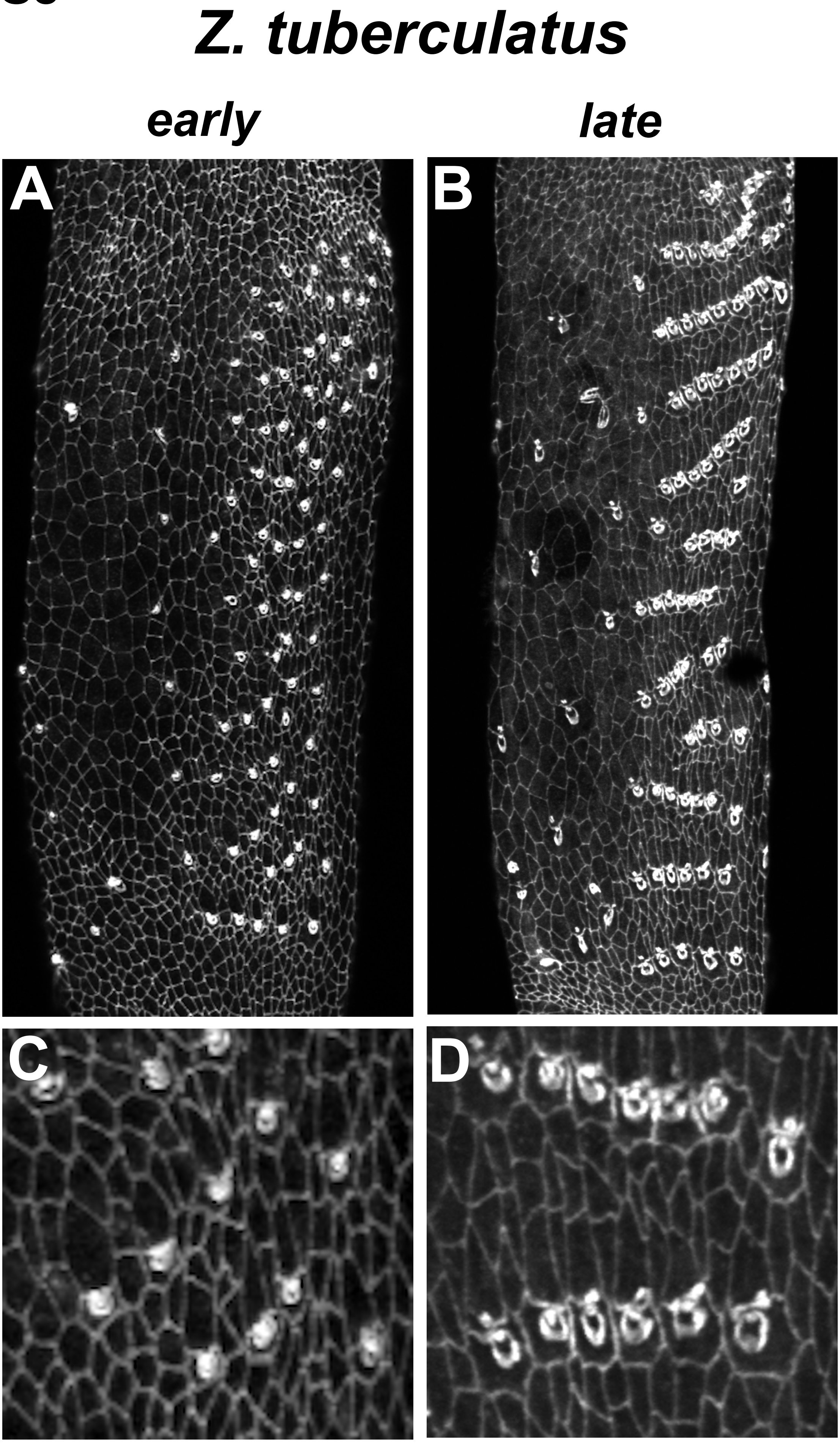

Supplemental Table S1. Sequence data used in the 8-locus phylogenetic analysis.

| | acon | glyp | eno | Pepck | Pgm | Amyrel | Gpdh | Ddc | Donor of live strain | Donor of fixed flies | Donor of unpublished genome | Reference for published genome |
|---|---|---|---|---|---|---|---|---|---|---|---|---|
| *D. pruinosa* | Accession # TBD | Accession # TBD | Accession # TBD | Accession # TBD | Accession # TBD | AY756177.1 | Accession # TBD | Accession # TBD | Jean-R. David | | | |
| *D. repletoides* | Accession # TBD | Accession # TBD | Accession # TBD | Accession # TBD | Accession # TBD | AY736500 | Accession # TBD | AF324978.1 | Ehime stock center | | | |
| *D. sternopleuralis* | Accession # TBD | Accession # TBD | Accession # TBD | Accession # TBD | Accession # TBD | AY736505.1 | AB932709.1 | Accession # TBD | | Masayoshi Watada | | |
| *D. trisetosa* | Accession # TBD | Accession # TBD | Accession # TBD | Accession # TBD | Accession # TBD | - - | Accession # TBD | Accession # TBD | | Masayoshi Watada | | |
| *D. curviceps* | Accession # TBD | Accession # TBD | Accession # TBD | Accession # TBD | Accession # TBD | Accession # TBD | Accession # TBD | Accession # TBD | Ehime stock center | | | |
| *D. quadrilineata* | Accession # TBD | Accession # TBD | Accession # TBD | Accession # TBD | Accession # TBD | MG684385.1 | AB261153.1 | MG684391.1 | Ehime stock center | | | |
| *D. annulipes* | Accession # TBD | Accession # TBD | Accession # TBD | Accession # TBD | Accession # TBD | - - | AB261152.1 | Accession # TBD | Masayoshi Watada | | | |
| *D. immigrans* | manual blast | manual blast | manual blast | manual blast | manual blast | AF491632.1 | AB261142.1 | AF293738.1 | | | Danny Miller | |
| *D. rubida* | Accession # TBD | Accession # TBD | Accession # TBD | Accession # TBD | Accession # TBD | AY736502.1 | MG684407.1 | MG684393.1 | Ehime stock center | | | |
| *D. albomicans* | manual blast | manual blast | manual blast | manual blast | manual blast | AF462595.1 | LC058572.1 | manual blast | | | | doi:10.1186/1471-2164-13-109 |
| *D. brachytarsa* | Accession # TBD | Accession # TBD | Accession # TBD | Accession # TBD | Accession # TBD | - - | Accession # TBD | Accession # TBD | | Stephane Prigent | | |
| *Z. indianus* | manual blast | manual blast | manual blast | manual blast | manual blast | EF458322.1 | manual blast | manual blast | | | Daniel Matute and Aaron Comeault | |
| *Z. tuberculatus* | manual blast | manual blast | manual blast | manual blast | manual blast | AY736524.1 | manual blast | manual blast | | | Daniel Matute and Aaron Comeault | |
| *Z. nigranus* | manual blast | manual blast | manual blast | manual blast | manual blast | EF458333.1 | manual blast | manual blast | | | Daniel Matute and Aaron Comeault | |
| *D. virilis* | XM_002052865.2 | XM_002051705.2 | XM_002052206.2 | XM_002049310.2 | XM_002046502.2 | XM_002048980.2 | D10697.1 | AF293749.1 | | | | |
| *D. americana* | manual blast | manual blast | manual blast | manual blast | manual blast | manual blast | EF137110.1 | AY165520.1 | | | | http://cracs.fc.up.pt/~nf/dame/index.html |
| *D. mojavensis* | XM_002003551.2 | XM_002001833.2 | XM_002002193.2 | XM_002006650.2 | XM_002011925.2 | genome blast | XM_002002668.2 | genome blast | | | | http://dx.doi.org/10.1038/nature06341 |
| *D. hydei* | XM_023314803.1 | XM_023319596.1 | XM_023310563.1 | XM_023307578.1 | XM_023311391.1 | AY733042.1 | L41650.1 | AF293737.1 | | | | |
| *D. grimshawi* | XM_001988581.1 | XM_001988878.1 | XM_001996423.1 | XM_001995058.1 | XM_001984373.1 | XM_001995261.1 | JN815819.1 | XM_001992779.1 | | | | |
| *D. busckii* | XM_017999532.1 | XM_017998597.1 | XM_017996761.1 | XM_017983203.1 | XM_017988961.1 | XM_017982827.1 | XM_017999954.1 | AF293733.1 | | | | |
| *D. nannoptera* | manual blast | manual blast | manual blast | manual blast | manual blast | manual blast | manual blast | manual blast | | | Virginie Orgogozo and Michael Lang | |
| *S. flava* | manual blast | manual blast | manual blast | manual blast | manual blast | - - | manual blast | manual blast | | | Noah Whiteman and Andy Gloss | |
| *S. pallida* | manual blast | manual blast | manual blast | manual blast | manual blast | manual blast | manual blast | manual blast | | | Noah Whiteman and Andy Gloss | |
| *D. melanogaster* | NM_001299190.1 | AF073177.1 | NM_001169382.2 | NM_001299647.1 | NM_079936.3 | NM_057914.4 | NM_057217.4 | genome blast | | | | http://dx.doi.org/10.1038/nature06341 |
| *D. pseudoobscura* | XM_001357457.3 | XM_001356612.3 | AY754465.1 | XM_001360906.3 | XM_001352495.3 | XM_001362009.3 | U47885.1 | AF293746.1 | | | | |
| *D. willistoni* | XM_002069248.2 | XM_002064779.3 | XM_002074300.3 | XM_002068739.3 | genome blast | AF039560.1 | L41248.1 | AF293750.1 | | | | http://dx.doi.org/10.1038/nature06341 |

Live flies were used whenever possible. If no live strain was available, fixed specimens were used instead. For sequences obtained from NCBI or generated for this study, Genbank accession numbers are provided. For unpublished or unannotated genomes where no gene models were available, sequences were extracted from fasta files of draft genome assemblies using stand-alone BLAST (v2.2.23).

Supplemental Table S2. Primers used to amplify gDNA sequences for the 8-locus phylogenetic analysis

| primer name | Sequence (5'->3') |
|---|---|
| acon F688 | GAGCTsAAGTGCCCCAAGGTGATTG |
| acon F869 | TCTGCAACATGGGTGCTGAGATCGG |
| acon R1763 | GTCAGATCCTkGCCGTTCCACTTGT |
| acon R1815 | GATGTGGTCrGTGGTGCACTTkCCC |
| eno F225 | CCGTCArATCTACGACTCCCGTGG |
| eno F426 | CGAGsTGATCAAGGCCAATCTGGA |
| eno R1247 | CCGATCTGGTTvACCTTCAGsAGC |
| eno R1433 | ATCTGGTTGTAyTTGGCCAGACGC |
| glyp F258 | sTCGCTGGAGTACTACATGGGTCGCTC |
| glyp F332 | AGGCCATGTACCAGmTkGGTCTGGACA |
| glyp R1255 | ACTTCTTCTTCACATTCTCCATrTGCA |
| glyp R1320 | GAyGCGCTTmTCGCCATCCTCCTCCACC |
| Pepck F712 | TGyGATCCcGAACGCACCATCATC |
| Pepck F950 | CCAAyCTGGCCATGyTGAAcCCCT |
| Pepck R1691 | ATCCAnTCgAGCACrCGgGAGTTC |
| Pepck R1875 | CACCTGsGACTCGAAGTAGkTGCG |
| Gpdh F | GTTCTAGATCTGGTTGAGGCTGCCAAGAA |
| Gpdh R | ACATATGCTCTAGATGATTGCGTATGCA |
| Ddc F | AGNCCCAAGTTNCATGCCTANNTTCC |

| | |
|---|---|
| Ddc R | ACCCAGCTGGGNTCCTTNAGCCACAT |
| Pgm F377 | TCGGCATCAAGTTCAACTGyGAGA |
| Pgm F575 | ATsTGCGyCwCATGGArGAAATCT |
| Pgm R1421 | CAAGGTCsAAGGAGGCsGACAACTT |
| Pgm R1432 | CyGTGTAGCTrAAGTTGTCsGCCT |

Supplemental Table S4. Prior distribution assumptions for model parameters used for ancestral character reconstruction.

* For all models, transition rate matrix is F81 model which is parameterized by the stationary frequency of the states; q ~ Dirichlet($\alpha_i,...,\alpha_K$) where K is number of states.

| | | | |
|---|---|---|---|
| MK unequal rates | Clock rate<br>Root frequency | r<br>$\pi$ | ~ Exponential(10)<br>= [0.5, 0.5] |
| RLC | Root frequency<br>Expected number of rate shifts<br>Probability of no rate shift on branch i<br>log Mean branch rate<br>Standard deviation of branch rates<br>Rate shift factor on branch i | $\pi$<br>n<br>$\rho_i$<br>$\log_{10} \mu$<br>$\sigma$<br>$\psi_i$ | = [0.5, 0.5]<br>= 2<br>= n/num_branches = 0.995<br>~ Uniform(-5, 2)<br>~ Exponential(1/0.587)<br>~ $\rho_i$ + (1 - $\rho_i$)logNormal($-\sigma^2/2$, $\sigma$) |
| Hidden rates class model | Root frequency<br>Transiton rates<br>log mean branch rate | $\pi_i$<br>q<br>$\log_{10} \mu$ | = 0.25 $\forall$ i $\in$ {1,...,4}<br>~ Dirichlet(1,1,1,1)<br>~ Uniform(-10,2) |
| Threshold model | Root frequency<br>Transition rates<br>log mean branch rate | $\pi_i$<br>q<br>$\log_{10} \mu$ | = 0.1 $\forall$ i $\in$ {1,2,...,10}<br>~ Dirichlet(⅓, ⅓, …, ⅓)<br>~ Uniform(-10,2) |
| Approximate dollo | Root frequency<br>Transition rates<br>log mean branch rate | $\pi$<br>q<br>$\log_{10} \mu$ | = [0.5, 0.5]<br>~ Dirichlet(1000, 1)<br>~ Uniform(-20, 2) |
